# Atomic structure and plasticity of the CTX-MthK complex investigated by cryo-EM, NMR, and MD simulations

**DOI:** 10.64898/2026.02.20.706983

**Authors:** Denis Qoraj, Swantje Mohr, Yessenbek K. Aldakul, Thiemo Sprink, Carl Öster, Taoran Xiao, Peter Schmieder, Sascha Lange, Tillmann Utesch, Daniel Roderer, Shanshuang Chen, Han Sun, Adam Lange

## Abstract

Scorpion toxins block potassium channels, disrupting cellular excitability and causing symptoms such as pain, muscle spasms, or paralysis. Here, we use an integrated structural biology approach to uncover the binding mode of the scorpion toxin charybdotoxin (CTX) to the MthK channel, a model system for human large-conductance potassium (BK) channels. Cryo-EM defines the overall architecture of the MthK–CTX complex, while complementary solution- and solid-state NMR experiments identify key binding residues and show that toxin engagement alters the selectivity filter (SF) ion configuration without rearranging the filter itself. NMR and MD simulations further reveal an anchoring lysine residue stably inserted into the SF, while other contacts undergo fast NMR timescale dynamics. Together, these findings explain how CTX-like toxins maintain exceptionally high affinity while tolerating binding across multiple K^+^ channel subtypes, paving the way for site-specific extracellular modulation.

## Introduction

Calcium (Ca^2+^)-gated potassium (K^+^) efflux is essential for numerous physiological processes, including muscle contraction, blood pressure regulation, and neuronal function, and its dysregulation is associated with a range of diseases and disorders^1,2^. The human large conductance Ca^2+^-gated K^+^ channel (also called MaxiK, BK channel) plays a central role in these processes and is distinguished by its unusually high conductance compared to other K^+^ channels^3,4^. BK channel activity is regulated not only by intracellular Ca^2+^ concentration, but also by membrane depolarization, enabling it to integrate both chemical and electrical stimuli^5,6^. Because structural characterization of large eukaryotic membrane proteins often requires complex expression, purification, and stabilization strategies, key mechanistic features of BK channels are frequently investigated using prokaryotic model systems. The Ca^2+^-gated K^+^ channel MthK, originally identified in archaea, represents one such well-established model protein^7^. Recent cryo-EM structures of MthK have revealed the atomic basis of the long-recognized N-type inactivation mechanism, which follows a “ball-and-chain” model^8^. Like BK channels, MthK is activated by intracellular Ca^2+^ through cytosolic calcium-binding domains (RCK domains) that assemble into a gating ring. In contrast, however, MthK lacks the voltage-sensing domains present in BK channels^7,9^, making it a simplified system for dissecting Ca^2+^-dependent gating mechanisms in isolation from voltage coupling.

The selectivity filter (SF) of MthK, located within the transmembrane domain and formed by the highly conserved TVGYG motif, constitutes the structural core of K^+^ selectivity. The backbone carbonyl oxygens of the filter coordinate four contiguous potassium-binding sites, thereby enabling highly selective and efficient conduction of K^+^ ions. Previous computational studies of MthK have suggested that K^+^ permeation proceeds via a direct knock-on mechanism, in which ions traverse the filter without intercalating water molecules within the pore^10–12^. Although the structures, gating mechanisms, and ligand interactions of MthK and BK channels have been extensively investigated, fundamental questions remain regarding the conformational dynamics of the SF and the ion conduction pathway, particularly under modulation by gating modulators and inhibitors, such as neurotoxins. One prominent example is charybdotoxin (CTX), a scorpion-derived peptide from the scorpion *Leiurus quinquestriatus*. Identified in 1985 for its potent inhibitory effect on Ca^2+^-activated K^+^ channels in mammalian skeletal muscle^13^, CTX subsequently became a key tool for mapping the pore region and establishing the tetrameric architecture of K^+^ channels before the first atomic structure of KscA was solved in 1998^14–16^. Similar to other pore-blocking toxins, CTX is a small peptide (∼4.3 kDa) stabilized by three disulfide bridges^17^. Furthermore, CTX and other toxins share a highly conserved lysine residue that protrudes into the SF of potassium channels, thereby physically occluding the ion conduction path^18^. Despite their relatively rigid fold and a well-defined binding motif, CTX and related toxins exhibit broad channel promiscuity, blocking multiple K^+^ channel families, including BK channels, *Shaker*, and MthK^7,13,19^.

Although numerous electrophysiological, mutational, computational, and structural studies^20–25^ have investigated neurotoxin binding to potassium channels, high-resolution structural information on intact channel-toxin complexes remains limited. Structural characterization of these assemblies is intrinsically challenging due to the symmetry-breaking nature of toxin binding and the small molecular size of peptides. These factors often result in poorly resolved or uninterpretable electron density maps in both X-ray crystallography and cryo-EM, caused by particle heterogeneity and averaging effects. To partially overcome these limitations, modified toxins have often been employed, for example, heavy-atom derivatives in X-ray crystallographic studies or fusion constructs to larger proteins in cryo-EM analyses^26,27^. However, such strategies reduce the number of available structures representing native neurotoxins bound to their physiological ion channel targets. Consequently, many mechanistic insights have relied on NMR studies of truncated channel constructs in detergent^24,28^ or in proteoliposomes^29^, frequently complemented by computational techniques^25,30^. Recent advances in detector technology, data processing algorithms, and cryo-EM software have begun to mitigate these challenges. Notably, the structure of the ∼7 kDa α-Dendrotoxin bound to Kv1.2 (∼242 kDa) was solved using cryo-EM^31^.

In the present study, we integrate the complementary strengths of cryo-EM, solution-state, and solid-state NMR to resolve the atomic-level binding mechanism of the small neurotoxin CTX to MthK, further supported by molecular dynamics (MD) simulations. This integrative structural approach enables us to define key interaction determinants governing MthK-CTX recognition and to showcase the structural plasticity of toxin binding to ion channels. Additionally, our data suggests that toxin binding induces a rearrangement of ion occupancy within the SF, while the filter itself remains in a non-collapsed and stable conformation.

## Results

### Overall architecture of the MthK-CTX complex

The MthK-CTX complex was first investigated using cryo-EM. For this purpose, MthK was reconstituted into nanodiscs composed of a 3:1 mixture of POPE:POPG, following the experimental strategy established by Nimigean and coworkers^8,32^, to enable direct comparison with previous structural studies of MthK. To form the MthK-CTX complex, CTX was added to MthK-containing nanodiscs prior to vitrification. Neither EDTA nor Ca^2+^ was included in the sample before analysis. First, a total of 106,517 particles were refined in cryoSPARC v4.6.2^33,34^, with C4 symmetry imposed, yielding a reconstruction at 3.2 Å resolution (SI Fig. 2). The resulting density map reveals MthK in a closed conformation, characterized by TM2 helices forming a bundle crossing at the intracellular side (Fig. 1A). Additionally, tightly bound lipids can be observed with their acyl-chains penetrating the hydrophobic gate through fenestrations, consistent with closed-state MthK structures^32^ (Fig. 1A, orange color). Additional density is observed at the extracellular pore entrance, resembling the symmetry-averaged CTX (Fig. 1A, green color). The high resolution of the map permits detailed analysis of ion occupancy in the SF, which is discussed more in details in a later section.

**Fig. 1:**
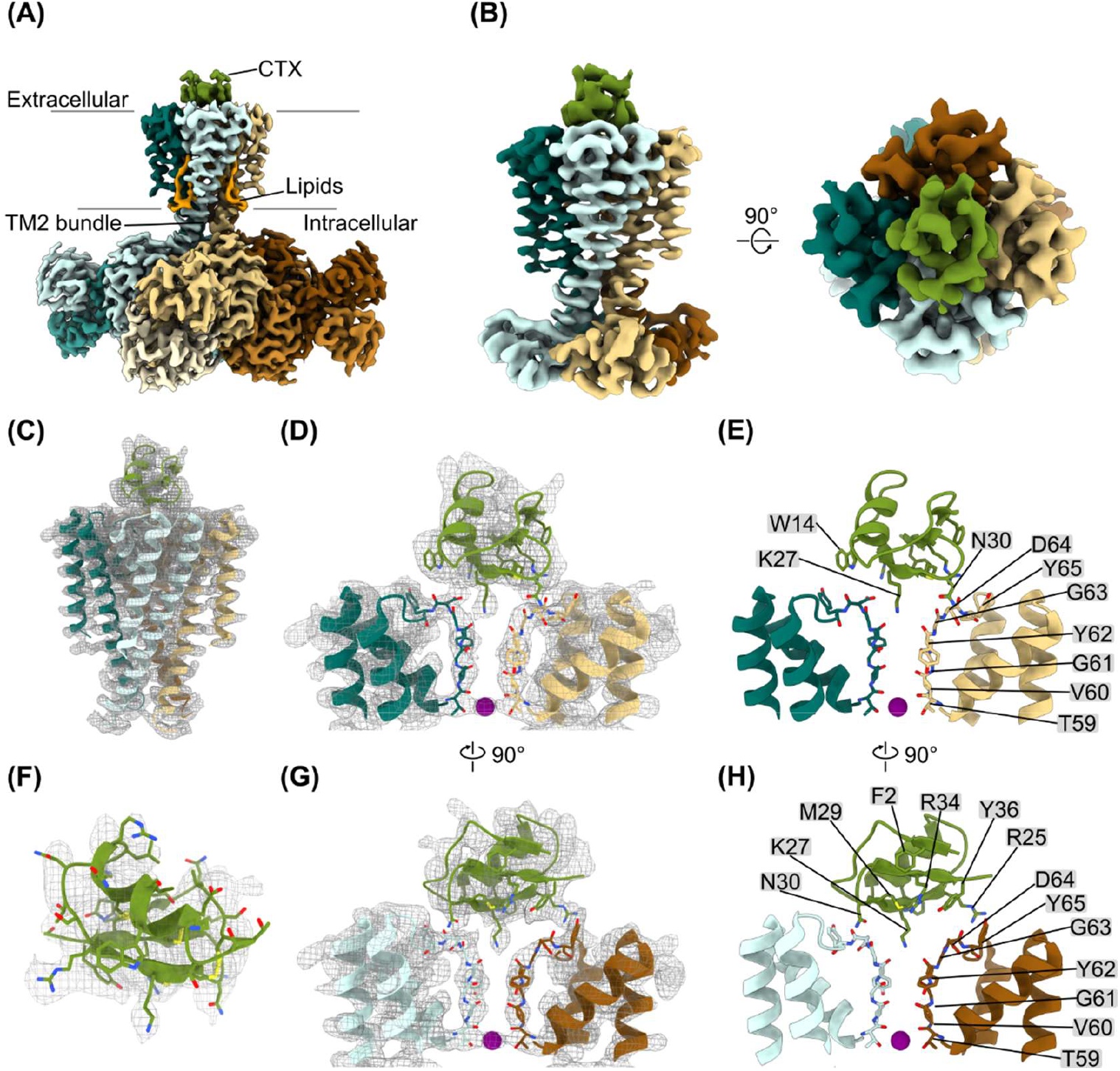
Cryo-EM analysis of the MthK-CTX complex. (A) Cryo-EM reconstruction of the full-length MthK-CTX complex obtained with imposed C4 symmetry to achieve the highest possible resolution. The density resulting from CTX is shown in green, while lipid densities are shown in orange. Each subunit is shown in a different color. (B) Particle-subtracted, asymmetric reconstruction of the MthK-CTX complex at an overall resolution of 4.1 Å. (C) Overall fit of the atomic model of MthK-CTX into the cryo-EM density map (EMDB: EMD-56657, PDB ID: 28NO). The map is shown as a gray mesh, and each MthK subunit is colored individually. (D-E, G-H) Close-up views of the MthK-CTX interface. CTX residues with well-resolved side-chain densities are highlighted, along the SF of MthK. For clarity, only two subunits of MthK are shown. (F) Fit of the CTX atomic model (green) into the corresponding cryo-EM density map (gray mesh). Additional examples in SI Fig.1. The complex structure was determined using Relion-5 and cryoSPARC v4.6.2.

Because CTX (∼4.3 kDa) is small relative to the homotetrameric MthK (∼250 kDa), reconstruction of the whole complex leads to substantial averaging of the toxin density due to symmetry mismatch. To enable reconstruction of an asymmetric map resolving CTX, we employed a workflow combining particle subtraction, symmetry expansion, and focused 3D classification in Relion-5 (details see Materials and Methods and SI Fig. 3). First, the gating ring of MthK, whose large mass dominates the particle alignment, was removed from raw particle images by particle subtraction. Symmetry expansion was then applied to generate multiple particle images from each original, matching all four-fold symmetric positions. This expanded particle set was subjected to focused 3D classification without alignment, minimizing overfitting and enabling the use of a small mask restricted to the MthK-CTX interaction region. To guide the 3D classification, we generated an Alphafold3^35^ model of the MthK-CTX complex and converted it into a density map that was low-pass filtered to 8 Å. The map was rotated in 90° increments around the symmetry axis to represent all possible toxin-binding orientations, following the approach described by Wu and coworkers in 2024^31^. These four maps served as templates for the 3D classification, yielding four classes corresponding to four distinct CTX orientations. One of these classes was selected for further refinement in Relion-5.

The resulting particle-subtracted, asymmetric structure of the MthK-CTX complex reached an overall resolution of 4.1 Å (Fig. 1B), with locally improved resolution at the protein-protein interface (∼3.8-3.9 Å, SI Fig. 1). In this region, the density is well defined, enabling confident modeling of several interacting residues and the interaction interface. The derived atomic structure shows CTX tightly bound to the extracellular side of MthK with the side-chain of K27 inserted into the SF at the S0 binding site, where it is coordinated by the backbone carboxyl oxygens of Y62 (Fig. 1C-D). Additional CTX residues at the interface, including W14, R25, N30, and Y36, also show visible side-chain features in the cryo-EM density (Fig. 1D-H). Within the resolution limits of the map, the SF of MthK (T59-G63) adopts a non-collapsed conformation. The density is continuous and consistent with the canonical conductive SF architecture, with no features indicative of filter collapse (Fig. 1D-E, G-H).

### Observing CTX chemical shifts in the absence and presence of MthK-PD via solution- and solid-state NMR

To further delineate the binding interface and probe the conformational dynamics of the MthK-CTX complex at atomic resolution, we performed complementary solution-state and solid-state NMR experiments on CTX, MthK, and the assembled complex^36,37^. The resulting data were analyzed in the structural framework provided by the cryo-EM structure to provide a comprehensive description of toxin engagement.

We first characterized ^15^N-labeled CTX in its free, soluble form using solution-state NMR. Accordingly, a set of 2D- and 3D-NMR spectra, namely 2D ^15^N-HSQC, 3D ^15^N-filtered NOESY, and TOCSY of the ^15^N-labeled toxin, were acquired. Most of the resonances of backbone N and HN atoms, as well as several side-chain atoms, could be assigned (Fig. 2A, yellow spectrum and SI Tab. S2), and - compared to previous assignments -, align well with the published data^20^. Only a single residue, M29 at the C-terminal end of the peptide, exhibited peak doubling, suggesting minor local conformational heterogeneity. This, together with the fact that our assignment fits very well with previously published data (BMRB ID: 114), indicates that the toxin adopts a correctly folded and well-defined structure in solution. This conclusion is further supported by torsion angle predictions derived from the assigned chemical shifts. The predicted secondary structures, mapped on the available PDB structure (PDB ID: 2CRD), show strong agreement with only minor deviations (SI Fig. 4). Additionally, we showed that when adding Ca^2+^ to the buffer, which is beneficial for the solid-state measurements of the MthK channel (as previously observed by our group^12^), no detectable changes in the NMR spectra of the soluble protein appear, indicating that Ca^2+^ does not perturb the structure of the soluble toxin (SI Fig. 5).

**Fig. 2:**
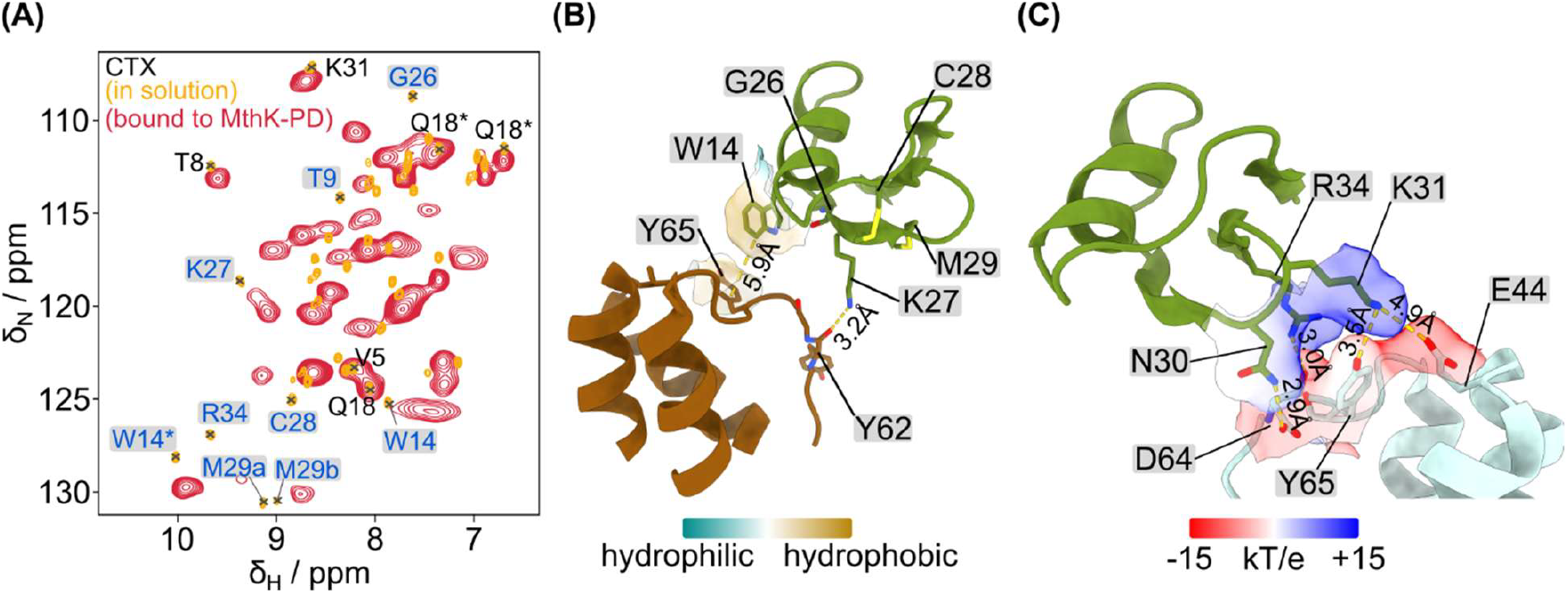
NMR analysis focused on CTX. (A) (H)NH solution-NMR spectrum of free CTX in yellow and solid-state NMR spectrum of bound CTX in red. Residues that show a clear chemical shift change are labeled in blue and highlighted with a gray box. Side chain signals are marked with an asterisk (*). (B) Detailed view of interacting residues between CTX and MthK, based on the determined cryo-EM structure and ssNMR experiments. Y65 (MthK) and W14 (CTX) show hydrophobic interactions, while Y62 (MthK) and K27 (CTX) form an electrostatic interaction. (C) Electrostatic interaction of the highly positively charged CTX with the negatively charged extracellular surface of MthK.

To investigate CTX in its MthK-bound state, proton-detected solid-state NMR experiments under fast magic-angle spinning (MAS) were performed. In these experiments, ^15^N-labeled CTX was added to the unlabeled MthK pore domain (MthK-PD, see Method section for further details) during reconstitution into proteoliposomes. Since CTX binds at the extracellular side of MthK, the use of a truncated pore-domain construct lacking the RCK domains reduces overall molecular size and improves spectral sensitivity and resolution. We recorded cross-polarization (CP) based (H)NH 2D spectra, which selectively detect rigid regions, as well as (H)NH spectra based on insensitive nuclei enhanced by polarization transfer (INEPT) that are sensitive to more dynamic residues. The CP based 2D spectra display well-resolved peaks (Fig. 2A, red spectrum), whereas no signals were observed in the INEPT-based experiments. This result indicates that CTX is immobilized upon binding and does not exhibit fast time-scale dynamics. Furthermore, the spectral dispersion and number of peaks are consistent with a well-folded toxin in the bound state, suggesting that CTX retains its structured conformation upon engagement with MthK.

Notably, by comparing the (H)NH-correlation spectra obtained from the free and the bound CTX (Fig. 2A), major differences could be identified. In addition to the expected line broadening in the solid-state spectrum, several resonances exhibit clear chemical shift perturbations, providing direct evidence of CTX interaction with MthK. However, direct transfer of resonance assignments from the solution-state NMR experiments of free CTX to the solid-state NMR spectra of the bound form is not straightforward. Nevertheless, several characteristic peaks in the less crowded regions of the spectra allow for detailed comparison: The peaks corresponding to backbone amide groups of V5, T8, Q18, and K31 overlap very well in the spectra (Fig. 2A). This indicates minimal changes in their local chemical environment upon binding. When mapping these residues onto the structure of CTX, with the exception of K31 (discussed later), they are predominantly located in loop regions or within the α-helical part of the toxin.

In contrast, the resonances of W14, G26, K27, C28, M29, and R34 exhibit notable changes or are absent in the spectrum of the bound CTX (Fig. 2A). While chemical shift perturbations can arise from changes in the chemical environment, the disappearance of resonances may be attributed to various factors: (i) large chemical shift changes can lead to peak overlap with other resonances. Since no complete assignment of the bound form of the toxin was achieved, it is difficult to distinguish between chemical shift changes and signal loss in the solid-state spectrum. ii) Changes in residue-specific dynamics may shift motion into a regime where CP transfer becomes inefficient, leading to attenuated or undetectable signals in the solid-state experiment. Irrespective of the exact reason, both chemical shift changes and missing peaks are both strong indications of the direct involvement of the respective residues in the interactions with the membrane-bound MthK channel. Structurally, these residues cluster within the β-sheet regions of the toxin, particularly around residue K27, which faces the extracellular pore entrance of MthK. Integrating these results with our findings from the cryo-EM investigation enables a mechanistic interpretation of these perturbations. The pronounced shift of K27 is consistent with its tight binding into the S0 K^+^ binding site of the SF (Fig. 2B). Perturbations observed for neighboring residues G26, C28, and M29 most likely arise from backbone adjustments induced by the tight anchoring of K27 within the SF, rather than direct side-chain interactions with MthK. Additionally, the ssNMR spectra of CTX show a clear shift of the W14 side-chain peak, consistent with a potential edge-to-face (t-shaped) stacking interaction with the phenol group of Y65 (Fig. 2A and B, distance: 5.9 Å).

In addition to K27, CTX contains several positively charged residues such as R25, R34, and K31, which, according to our cryo-EM structure, contribute to tight binding at the negatively charged extracellular pore surface of MthK. The ssNMR spectra show a clear chemical shift change for R34 in CTX, which interacts with the CO of D64 in MthK (Fig. 2A and C, distance 3.0 Å). In proximity, the NH_3_^+^ group of K31 (CTX) is located in a negatively charged pocket formed by the aromatic OH group of Y62 (MthK) and the carboxyl group of E44 (MthK, Fig. 2C), while N30 (CTX) forms a hydrogen bond with the carboxylate group of D64 (MthK) near the pore entrance (Fig. 2C, distance 3Å). The comparatively small chemical shift changes of K31 (CTX) are consistent with an interaction mediated primarily through its flexible side-chain ammonium group, whereas the backbone amide detected in the solid-state NMR spectrum is further away from the primary interaction site. Unfortunately, the side-chains of lysine residues were not unambiguously assignable in the solution-state spectrum, and consequently could not be evaluated in the solid-state spectrum.

### Solid-state NMR investigation of changes in MthK-PD caused by CTX binding

To obtain additional high resolution structural information on the complex from the perspective of MthK, we performed ^1^H-detected solid-state NMR on isotope labeled MthK. Specifically, spectra of 100% H_2_O back-exchanged ^2^H^13^C^15^N labeled MthK-PD with or without unlabeled CTX were compared. Based on resonance assignments of the free MthK-PD previously published from our lab^12^, MthK-PD in the presence of CTX was assigned.

Comparison of the spectra of MthK-PD with and without CTX reveals that the majority of chemical shift changes localize to the SF region or its vicinity (Fig. 3A and B). In particular, residues Y62 to D64 (MthK), some of which are present in two different conformations in the free channel, show a large shift of about 2 ppm in the nitrogen dimension (Fig. 3A and B). Additionally, only a single conformation was detected for Y62, likely attributed to the stabilization of one conformation relative to others. To assess whether these chemical shift perturbations arise from structural rearrangements of the respective amino acid residues, the Cα-chemical shifts were analyzed. Interestingly, apart from Y62, the changes are below 1 ppm, and the largest differences can instead be observed at residue T69, located in the extracellular loop (Fig. 3D). Together with the cryo-EM structure, which does not show significant structural changes in the SF upon CTX binding, these results suggest that the observed chemical shift changes primarily reflect modifications of the local electrostatic environment and ion occupancy within the filter upon toxin binding, rather than structural disruptions such as a collapsed SF.

**Fig. 3:**
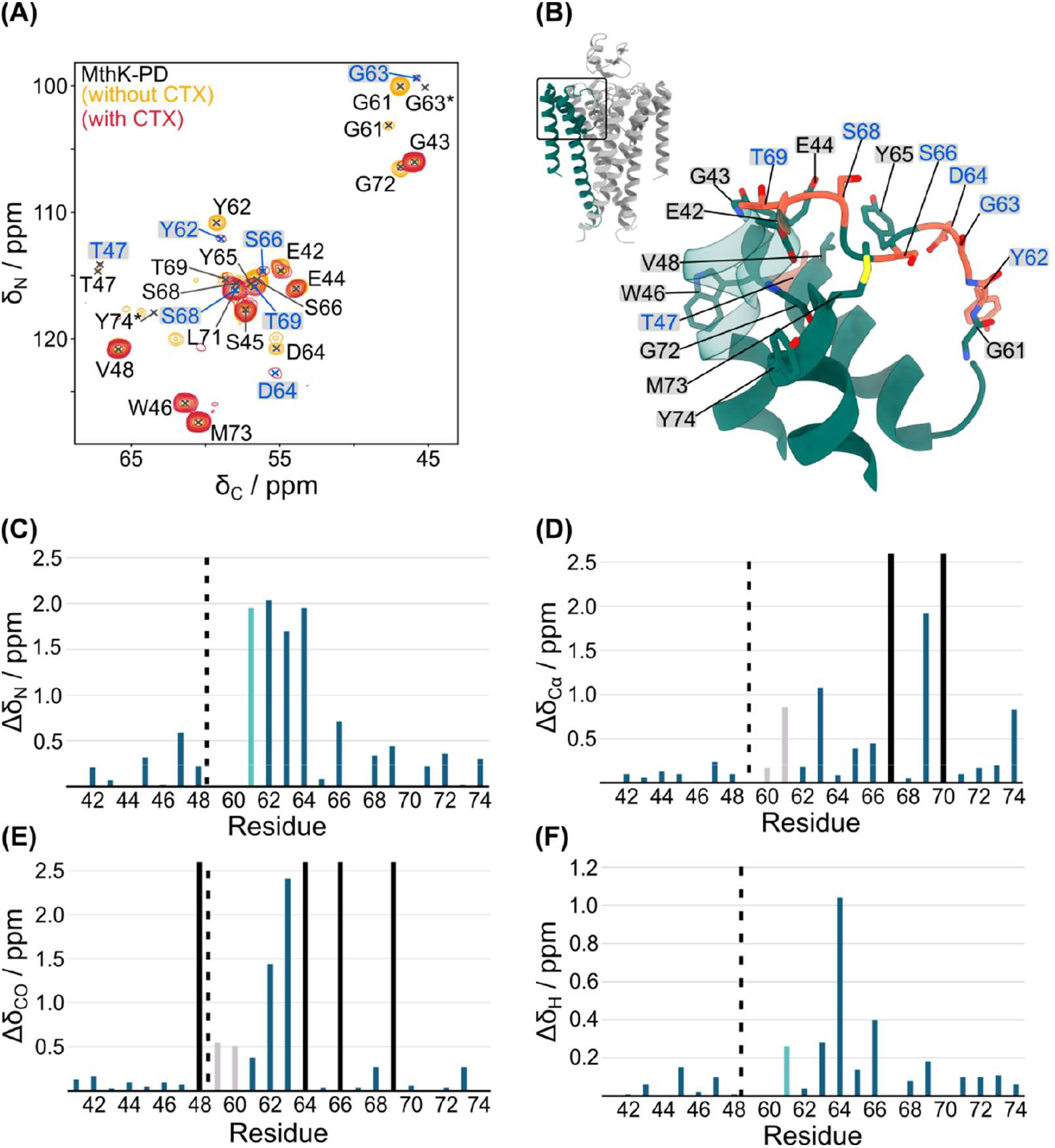
NMR analysis focused on MthK. (A) CαN projection of a (H)CαNH ssNMR experiment of MthK-PD with and without CTX. Residues showing chemical shift perturbations (CSPs) are labeled in blue and highlighted by a gray box. Resonances not visible in the projection, but in the 3D spectra, are marked with an asterisk (*). (B) Zoom-in on the MthK extracellular and SF regions of the MthK-CTX complex. All residues that could be assigned through ssNMR experiments are shown as sticks. Residues showing chemical shift changes upon CTX binding are colored red and labeled in blue. (C-F) Individual CSPs of the backbone atoms from MthK-PD induced by CTX binding, calculated as described in the Method section. Residues not assigned in the CTX-bound sample are shown with black bars, chemical shifts assigned on the samples with washed-in CTX or ^15^NH_4_ ^+^-containing buffer are colored in light blue and gray, respectively.

G61 in MthK showed no detectable signals in the CTX-bound sample, which may reflect reduced water accessibility that limits the H/D exchange required for detection. To further investigate this effect, an additional MthK-PD sample was prepared in which CTX was introduced after reconstitution by washing the pelleted sample with CTX-containing buffer for three days to ensure complete binding. Interestingly, here a resonance belonging to G61, based on comparison to assignments of the free MthK-PD, could be detected in the H-N correlation spectrum (SI Fig. 6).

As all other residues showed no chemical shift differences compared to the sample in which CTX was added during dialysis, the observations are consistent with CTX altering the dynamics of the SF, thereby restricting H/D exchange of the backbone amide proton of G61, highlighting the tight binding of CTX.

The chemical shift changes of residues on the extracellular loops cluster on residues likely involved in electrostatic interactions with the toxin interface, as already discussed in the previous section. Here, the new chemical environment induced by the highly positively charged CTX surface generates chemical shift perturbations, especially visible for partially negatively charged or polar residues, including D64, S66, and T69 (Fig. 3C-F).

### MD simulations show dynamic binding of CTX to MthK and reveal the amplitude and time scale of this motion

To further characterize the dynamics and stability of CTX binding to MthK, all-atom MD simulations of the complex were performed. The starting model of CTX bound to MthK-PD was derived from the cryo-EM structure. Across all three runs of 500 ns MD simulations, CTX remained bound to MthK, with K27 of the toxin consistently inserted into the S0 site of the SF (Fig. 4A and C). Root-mean-square fluctuation (RMSF) analysis of CTX indicates an overall stable toxin conformation, reflected by low RMSF values of structurally constrained cysteines, i.e. C7 and C28, involved in disulfide bridges (Fig. 4B). While anchored via K27 at S0, CTX retains a certain degree of orientational flexibility, undergoing rotational motions in our simulations up to ∼20° relative to the channel axis (Fig. 4A). Interactions observed in the cryo-EM structures, such as between R34 (CTX) and D64 (MthK) or W14 (CTX) and Y65 (MthK), transiently break and reform during the simulations (Fig. 4C and SI Fig. 7), typically on a 1-100 ns timescale, reflecting dynamic stabilization of the interface.

**Fig. 4:**
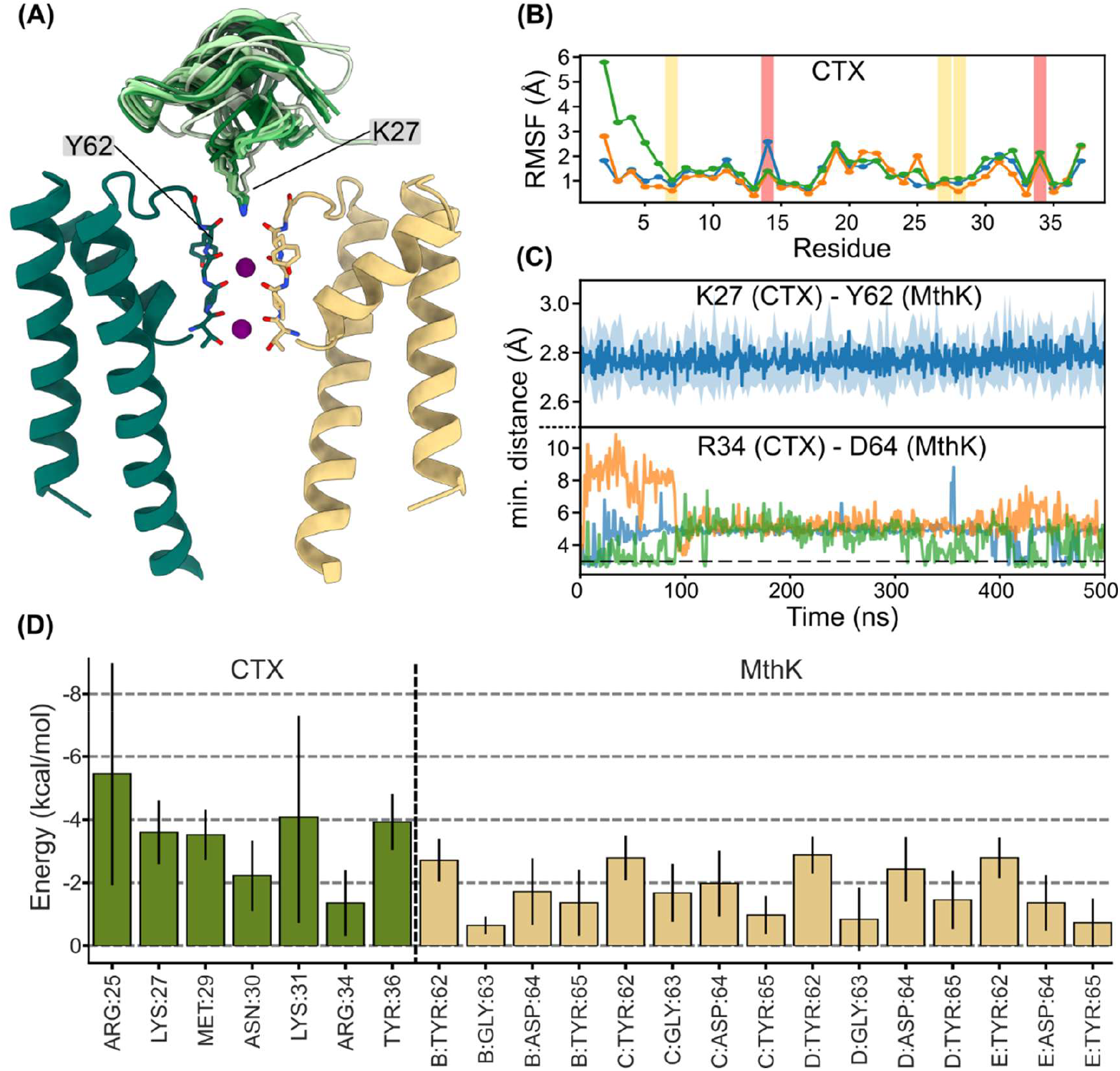
MD simulations of the MthK-CTX complex. (A) Snapshots from MD simulations, where the starting structure is based on the cryo-EM-derived MthK-CTX structure. The CTX conformations from every 50 ns are shown, colored from green to white. K27 is shown in stick representation and anchors the toxin to the SF of MthK. (B) RMSF values of CTX from three runs (different colors) of 500 ns MD simulations. Flexible (highlighted in red) and rigid (highlighted in yellow) residues mentioned in the text are marked with boxes. (C) Minimum distances between different residues during MD simulations. The average minimum distance between K27 (CTX) and Y62 (MthK) residues is shown as a solid blue line with a transparent blue shade showing the standard error of the mean (Data from three MD simulation runs). The minimum distances between R34 (CTX) and D64 (MthK) are shown separately, with each line corresponding to one MD run. (D) Binding free energy calculations with the Generalized Born model. Vertical lines indicate the standard deviation. Green bars show residues from CTX, while residues from MthK are shown in beige.

To quantify binding energetics, we estimated the binding free energy of the complex using MM/PBSA calculations (Fig. 4D)^38^. Results show that binding of CTX to MthK is energetically favorable. Residues contributing most strongly to binding localize to the interface of CTX and MthK, in agreement with cryo-EM and NMR results. On the channel side, Y62 residues within the SF interact strongly with K27 of CTX, emphasizing the tight binding of these residues. Due to the asymmetric binding mode, other MthK residues contribute variably to the overall binding energy. On the toxin side, in addition to K27, the adjacent residues M29 and N30 of CTX also contribute favorably to binding. R25 and K31 (CTX) also exhibit strong stabilizing interactions with MthK; however, consistent with the observed rotational flexibility of CTX, these interactions are dynamic and associated with relatively large free energy standard deviation in the free energy estimates.

### Ion configuration of MthK changes upon CTX binding

Solid-state NMR reveals pronounced chemical shift changes in the upper region of the SF. However, neither the cryo-EM map nor the NMR spectra support a collapsed or structurally altered SF, suggesting that the observed shifts, especially for the central SF residues, originate primarily from changes in ion configuration upon toxin binding. To investigate the ions in more detail, we compared the SF of the C4 reconstructed map of the full-length MthK-CTX complex, which reaches a higher resolution of 3.2 Å compared to the asymmetric reconstruction (4.1 Å), with published toxin-free MthK structures. Four discrete densities are visible in the SF of MthK (Fig. 5A). The density at S0 (green) corresponds to the ε-amino group of K27 from the bound CTX, while the other three densities (purple) are consistent with K^+^ ions bound in the SF (Fig. 5A-C). In contrast to the published free MthK channel (EMDB: 9405, PDB ID: 5BKI)^32^, where S1-S4 are occupied with K^+^ ions, the CTX binding alters the ion configuration of MthK (Fig. 5D-F). The S1 binding site is unoccupied and the density between S2 and S3 appears smeared. The reduced definition of the ion densities relative to the channel may indicate partial occupancy at these sites in the CTX-bound state. Since cryo-EM maps do not directly report dynamics but an average over all used particles in the dataset, we next directly probed ion binding using ssNMR with ^15^NH_4_^+^ ions as an NMR-visible alternative to K^+^ ^39–41^.

**Fig. 5:**
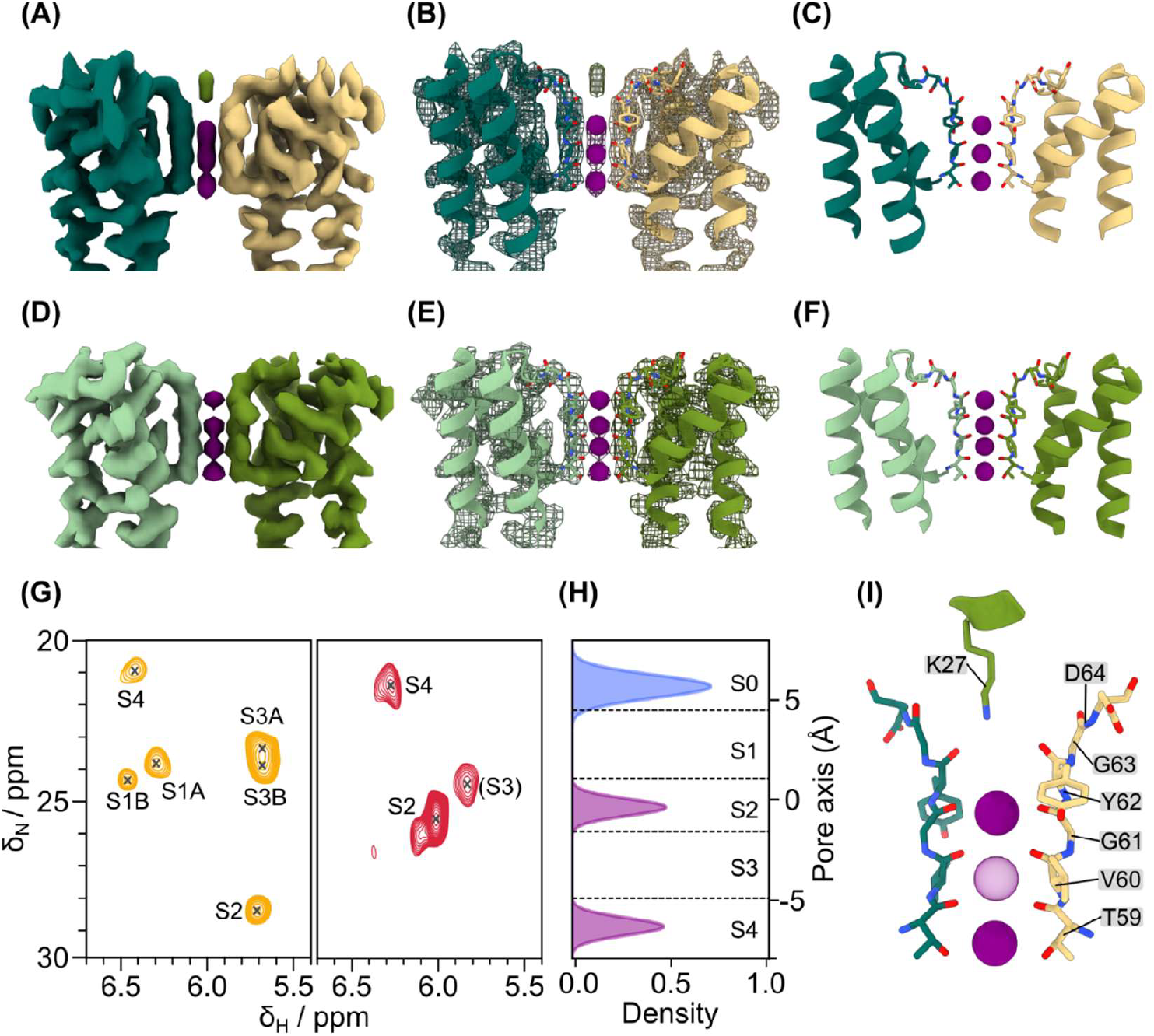
Ion occupation in CTX-bound MthK. (A-C) Full-length symmetric cryo-EM map of the MthK-CTX. The map and a relaxed model based on the published 5BKI^32^ structure is shown here. (D-F) The cryo-EM map and atomic structure of the published closed-state free MthK^32^. (G) (H)NH INEPT 2D spectrum of the bound ^15^NH_4_^+^ ions in MthK-PD without (yellow spectra, reproduced from^12^, and with CTX (red spectra). Ions in brackets could not be unambiguously assigned. (H) The occupancies of K27 (blue) and K^+^ (purple) in the SF binding sites during MD simulations. (I) Integrative model of the ion occupation in the MthK-CTX complex.

Replacing K^+^ ions in the MthK channel with ^15^NH_4_^+^ causes only minor changes in backbone chemical shifts for the SF residues^12,40^, enabling site-specific analysis. To examine the effect of the toxin, both free MthK-PD and the complex were studied with 100 mM of ^15^NH_4_^+^ instead of K^+^ ions. Analyzing the ammonium signals revealed significant differences between the two samples. In the (H)NH INEPT 2D fingerprint spectrum of the free MthK-PD, six signals were observed, whereas only four peaks were detected in the CTX-bound state (Fig. 5G). The six resonances in the free MthK-PD originate from multiple ion configurations in the SF, leading to multiple states for ^15^NH_4_^+^ ions bound to S1 and S3. To assign the ^15^NH_4_^+^ resonances seen in the complex to specific ion-binding sites within the SF, we used the approach described in our previous study with (H)CH experiments and 7 ms CP mixing^39,40^. Two signals exhibit contacts with Cα and CO atoms of SF residues, corresponding to the S2 and S4 binding sites (SI Fig. 8). The S2 signal shows higher intensity, suggesting higher occupancy. The most upfield-shifted peak in the proton dimension overlaps with the backbone HN signal of E44, making it difficult to identify H-C contacts. While it may represent the S3 binding site, supported by cross peaks to the CO of V60, an unambiguous assignment is not possible. Compared to the free channel, S4 shows only minor chemical shift changes, while S2 exhibits substantial perturbations in both proton and nitrogen dimensions. This likely results from the nearby K27 side-chain of CTX, causing a significant change in the local chemical environment. Together, the ssNMR data indicate displacement of the S1 ion and preferential occupancy of S2 and S4 in the CTX-bound state, while multiple configurations may still coexist.

To further evaluate ion occupancy, molecular dynamics simulations were performed with one or two K^+^ ions in the SF. Simulations with 2 K^+^ ions in the SF converged to a stable conformation with the ε-amino group of K27 in S0 and K^+^ ions in S2 and S4, even when initialized at S3/S4. This behavior may arise from electrostatic repulsion from K^+^ bound at S4 or the favorable binding of K^+^ at S2 over S3. To evaluate whether force-field electrostatics contribute to this preference, we performed additional MD simulations using modified ion parameters that introduce the polarization effect in a mean-field manner (SI Fig. 7C)^42^. These simulations, again, showed the translocation of K^+^ from S3 to S2, hinting at a preference of S2 in the CTX-bound state. MD simulations with one K^+^ ion in the SF match the observed preference of S2 and additionally resulted in a collapsed SF with water entering the pore (SI Fig. 7B-C). Overall, MD simulations agree with the ssNMR data and support the predominant occupation of S2 and S4, while ions likely exchange between nearby binding sites within the filter.

## Discussion

In this study, we solved the structure of the MthK-CTX complex using the wild-type toxin and an integrative approach combining cryo-EM, solution- and solid-state NMR, and MD simulations. With the overall binding position of the toxin matching the literature, key residues involved in the binding could be identified by cryo-EM and NMR data. On the channel side, no large structural changes could be observed from the cryo-EM structure, while chemical shift changes in NMR experiments were mostly seen for residues of the SF and the upper part of the pore at the protein-protein interface. These can be explained by a changed chemical environment upon toxin binding rather than large structural rearrangements. These observations are in line with previous crystallographic studies, suggesting a binding of the neurotoxin only to the upper surface of the K^+^ channel^26^. In a previous ssNMR investigation on the binding of KTX to the chimeric channel KcsA-Kv1.3, chemical shift changes, e.g., for Y78 Cα (KcsA-Kv1.3) - indicative of backbone torsional angle changes - as well as distance restraints involving Y78 recorded using NHHC experiments^29^ pointed towards structural changes in the upper part of the SF. An investigation of this system with the methods employed in the current study may be worthwhile, in particular, the direct detection of ions using the ammonium approach. However, differences observed may also depend on the specific system of interest, as the selectivity filter in different ion channels may respond differently upon toxin binding.

Residues shown to be important for the toxin binding in early mutation studies on similar K^+^ channels^23,43–45^, such as D64 and the upper SF residues G61 and Y62, were found to be in proximity to the toxin. They also exhibit chemical shift changes induced by the close surface of the binding partner. On the toxin, residues around the highly conserved K27, especially G26, C28 and M29, which are all by themselves conserved across multiple neurotoxins^44^, were found to interact in cryo-EM and NMR experiments, as suggested in multiple studies on CTX^21,46^ and related toxins^23,47^. Even though there is no clear interaction partner for M29 (CTX), the residue is in proximity to R34 (CTX) and Y36 (CTX) and could be involved in coordinating these residues. Additionally, residues interacting with the upper part of the channel pore, such as W14, N30, and R34 (all CTX) were found to interact in the cryo-EM map, also corroborating previous findings. Interestingly, W14 is important for the binding of CTX to BK channels, while mutations of this residue exhibit no effect on the binding to Shaker^21^. Although MthK, BK and Shaker each harbor hydrophobic residues adjacent to the SF, only Y65 in MthK and F283 in the BK channel display exposed aromatic rings. In contrast, the side-chain of M448 in Shaker is buried, hindering an interaction with W14^7,48,49^.

MD simulations revealed a dynamic behavior of the toxin while being bound to MthK. Although the overall fold is preserved, CTX undergoes some rotational and tilting motions on the extracellular surface of the channel within the limited simulation time scale, while K27 remains consistently inserted into the SF. This dynamic binding behavior firstly offers a plausible explanation for the absence of peak doublings in the solid-state NMR spectra of the channel, which would otherwise be expected if toxin binding rigidly disrupted the fourfold symmetry of MthK, and secondly could be a reason for the binding plasticity of CTX. Despite its high affinity for diverse K^+^ channels, the toxin does not adopt a single rigid binding pose. Instead, flexible side chains engage in transient interactions with different channel residues, with contacts continuously forming and breaking. The side-chain of K27 acts as an anchor, keeping the toxin bound to the SF. This gives CTX the flexibility to bind to various K^+^ channels with minor changes to their extracellular surface structure. Additionally, this dynamic binding mode may help explain the challenges associated with the structural determination of channel-toxin complexes. In cryo-EM reconstructions, CTX density is already weakened by symmetry averaging due to its small size. Additional small-amplitude rotational motions further blur the density, making high-resolution reconstruction of the toxin particularly demanding.

Upon CTX binding, the ion configuration of MthK is altered, supported by both cryo-EM and ssNMR data. The symmetric cryo-EM map reveals four densities within the SF: the K27 side-chain of CTX occupies the S0 site, while S2, S3 and S4 are occupied by K^+^ ions. The diffuse densities observed at S2 and S3 suggest a partial occupancy or increased ion dynamics at these positions. Complementary ssNMR experiments with ^15^NH_4_ ^+^ ions clearly show bound ions in S2 and S4, and an additional resonance consistent with S3, although this assignment remains ambiguous and the peak is relatively weak, suggesting reduced occupancy. This would result in an altered pattern with S0 (by K27 from CTX), S2, and S4 occupied, while S1 and S3 stay empty, distributing the positive charges throughout the SF. A theoretical possibility would be that the filter contains only one ion that exchanges between binding sites, or that the sample consists of multiple configurational states in which the ion resides at S2 in some channels and at S4 in others. However, when simulating the complex with only one K^+^ ion placed in S2 or S3, results show preferential binding of K^+^ in S2 (SI, Fig. 6) and a collapsed SF. Interestingly, if K^+^ is placed only in S4 the SF also collapses, but the ion stays bound to S4 (SI Fig. 7). Nevertheless, since neither cryo-EM nor ssNMR experiments suggest a collapsed SF, this suggests that the SF most likely is occupied by at least two ions (i.e. S2 and S4), and that the ion at S2 possibly moves between S2 and S3.

In conclusion, we resolved the atomic structure of the MthK-CTX complex and revealed the structural plasticity underlying CTX binding to K^+^ channels. CTX can be conceptualized as a molecular “master key” capable of inhibiting diverse K^+^ channel architectures. While K27 functions as the central anchoring residue by inserting into the SF, residues with flexible side-chains, including W14, R34, and K31, adapt their interactions to accommodate variations in the extracellular surface of different K^+^ channels. These mechanistic insights provide a structural framework for the rational design of new inhibitors or engineered binding partners for K^+^ channels. Importantly, the integrative strategy employed here, combining cryo-EM, solution- and solid-state NMR, and MD simulations, demonstrates how high resolution structural information can be achieved in systems where any single method alone would be insufficient^50^. Beyond defining the toxin-channel interface, this approach further enables atomic-level investigation of ion-binding equilibria within the SF, a fundamental biophysical question that remains the subject of ongoing debate.

## Author Contributions

D.Q., T.X., and S.L. prepared cryo-EM and NMR samples; D.Q. and T.S. performed cryo-EM experiments; D.Q, T.S., D.R., S.C., and T.U. analyzed cryo-EM data; S.M. and P.S. performed and analyzed solution NMR experiments; S.M. and C.Ö. performed and analyzed solid-state NMR experiments; Y.K.A. and H.S. performed and analyzed MD simulations; A.L. designed the study and supervised the work. The manuscript was written by D.Q., S.M., and Y.K.A. with input from all authors and revised by H.S. and A.L. All authors have given approval to the final version of the manuscript.

## Notes

The authors declare no competing financial interest.

## Acknowledgments

We thank Prof. Misha Kudryashev for discussions. We thank the Core Facility for cryo-Electron Microscopy (CFcryo-EM) of the Charité—Universitätsmedizin Berlin for support in acquisition of the data. The CFcryo-EM was supported by the German Research Foundation (DFG) through grant No. INST 335/588-1 FUGG. This work was funded by the Leibniz-Forschungsinstitut für Molekulare Pharmakologie, the Leibniz Society within the Leibniz Collaborative Excellence funding program (Project title: Ion Selectivity and Conduction Mechanism of Cation Channels, Project number: K305/2020 - to A.L. and H.S.) and the Deutsche Forschungsgemeinschaft (DFG, German Research Foundation) – under Germany’s Excellence Strategy – EXC 2008/1 (UniSysCat) – 390540038 (to H.S. and A.L.).

## Data deposition

The cryo-EM map of the truncated asymmetric MthK-CTX complex has been deposited in the Electron Microscopy Data Bank (EMDB) under the accession code EMD-56657. Atomic coordinates for the MthK-CTX complex have been deposited in the Protein Data Bank under the accession code 28NO. Assigned chemical shifts obtained from solution NMR spectra on CTX were deposited in the BMRB under the ID 53511.

## Materials and Methods

### Expression and purification of MthK full-length and pore domain

The native MthK channel gene was synthesized and cloned in pQE60 (BioCat GmbH, Heidelberg, Germany) with a carboxy-terminal hexahistidine tag (His_6_-Tag), including a tobacco etch virus (TEV) cleavage site before the His_6_-Tag. The vector was transformed into XL1-blue cells (Agilent) and grown at 37 °C in Luria-Bertani (LB) medium supplemented with 100 µg ml^-1^ carbenicillin. At a cell density (OD_600_) of ∼0.8, the expression of MthK was induced by adding 400 µM isopropyl-β-D-thiogalactoside (IPTG), and cells were grown for 3-4 h at 37 °C. The cells were harvested via centrifugation at 4000 relative centrifugal force (rcf) at 4 °C. The pellet was resuspended in 10 mL lysis buffer (100 mM KCl, 50 mM Tris, pH 7.6 with 1 µM leupeptin/pepstatin (MERCK) and 0.17 mg ml-1 PMSF (Roche)) per gram wet cell mass. Before cell lysis, lysozyme and DNase were added to the resuspended cells. Cell lysis was performed with the LM10 microfluidizer (Microfluidics, USA) at 15,000 psi working pressure. Membrane proteins were solubilized by adding 2% (w/v) n-decyl-ß-D-maltopyranoside (DM, Glycon Germany) to the cell lysate and incubating for 3 h at room temperature. Insoluble cell debris was removed via centrifugation at 18.000 rcf for 45 min at room temperature. The supernatant was supplemented with 20 mM imidazole and incubated for 30 min with Co^2+^-loaded Talon Beads (TALON^®^ Superflow™, Cytiva, USA) equilibrated with equilibration buffer (100 mM KCl, 20 mM Tris-HCl, 0.2% (w/v) DM, pH 7.6) at room temperature. The beads were washed with 10 bed volumes (BV) wash buffer (100 mM KCl, 20 mM Tris-HCl, 20 mM imidazole, 0.2% (w/v) DM, pH 7.6) and MthK was eluted with 5 BV using 450 mM imidazole. Desalting was performed using a HiPrep Desalting column equilibrated with 100 mM KCl, 20 mM Tris-HCl, 0.2% (w/v) DM, pH 7.6. Subsequently, the His_6_-Tag was cleaved overnight at room temperature by adding 0.2 mg His-tagged TEV protease per 1 mg of purified protein. A reverse-IMAC was performed, and the protein was concentrated to 10 mg ml^-1^ using 50,000 MWCO Amicon concentrators (Millipore).

Perdeuterated, ^13^C- and ^15^N-labeled MthK was expressed and purified as described previously^12^. In short, a codon-optimized DNA sequence encoding full-length carboxy-terminal His_6_-tagged MthK was synthesized by GeneArt Life Technologies (Germany), cloned into the pET21a vector, and transformed into *E. coli* BL21(DE3)pLysS. Bacterial cultures were grown in perdeuterated M9 minimal medium with ^15^ND_4_Cl (Cambridge Isotope Laboratories, USA) and ^2^H,^13^C labeled glucose (Cambridge Isotope Laboratories, USA) as the sole nitrogen and carbon sources, following a previously published deuterium adaptation protocol^51,52^. The protein was overexpressed for 16 hours at 25 °C after IPTG induction (0.5 mM) at an OD_600_ of 0.8. Cells were harvested, resuspended in lysis buffer (20 mM Tris-HCl, 100 mM KCl, 1.4 mM ß-mercaptoethanol, pH 8.0), and lysed using an LM10 microfluidizer (Microfluidics, USA) at 15,000 psi working pressure. The cell lysate was centrifuged at 18.000 rcf for 45 min at 4 °C, and the supernatant was incubated with 2% DM (w/v) for 2 h at 4 °C. The solubilized protein was purified by cobalt-based gravity flow affinity chromatography (TALON^®^ Superflow™, Cytiva, USA) with 0.2% DM (w/v) in wash (20 mM Tris, 100 mM KCl, 10 mM imidazole, 1.4 mM ß-mercaptoethanol, pH 8.0) and elution buffer (20 mM Tris, 100 mM KCl, 500 mM imidazole, 1.4 mM ß-mercaptoethanol, pH 8.0). Purified full-length MthK was digested with bovine pancreatic trypsin (Sigma-Aldrich) for 2 h at room temperature, and the reaction was stopped by the addition of trypsin inhibitor type II (Sigma-Aldrich).

Isotope labeled, trypsin digested MthK pore domain (PD) was mixed with asolectin (Sigma-Aldrich) in 5% DM (w/v) at a lipid to protein ratio of 1:1 (w/w). Detergent was removed by dialysis against sample buffer (20 mM Tris, 100 mM KCl or 100 mM ^15^NH_4_Cl, 10 mM CaCl_2_, and1.4 mM ß-mercaptoethanol, pH 8.0) over the course of 10 days and buffer exchanges every second day. For the samples with added CTX, the toxin (labeled or unlabeled) was added in a 4:1 molar ratio (CTX : MthK tetramer) to the protein in detergent before dialysis. The sample was centrifuged for 2 h at 4 °C and 100,000 rcf, and the pellet was used to fill MAS rotors by centrifugation in a benchtop centrifuge at 10,000 rcf.

### Unlabeled and labeled CTX expression and purification

The codon-optimized DNA coding for the fusion protein His_6_-SUMO-CTX was synthesized and cloned in pET28a (BioCat GmbH, Heidelberg, Germany). Unlabeled protein expression was initiated by inoculating 5 ml of LB medium with a single colony of a freshly transformed *E. coli* HMS174 (DE3), while ^15^N-labeled protein was expressed in M9 minimal medium with ^15^NH_4_Cl (Cambridge Isotope Laboratories, USA) as the sole nitrogen source. Following a 2.3 h incubation period at 37 °C, the preculture was diluted into 200 ml of fresh LB medium. The second preculture was incubated overnight at 30 °C and used to inoculate 2 L of fresh LB or M9 minimal medium to an OD of 0.2. Following an incubation period at 37 °C until an OD_600_ of 0.8–1.0 was reached, induction of protein expression was initiated by the addition of 1 mM IPTG. Bacteria were harvested after 16 hours by centrifugation at 5000 rcf and stored at -80 °C. The frozen cell pellets were resuspended in 100 ml of lysis buffer (6 M guanidine hydrochloride (GuHCl), 10 mM imidazole, pH 8.0) and disrupted by sonication (40% power, with three intervals of 5 min each). Subsequently, the lysate was subjected to centrifugation at 20,000 rcf for one hour, after which the supernatant was diluted with phosphate-buffered saline (PBS) containing 10 mM imidazole to a volume of 500 ml. This cleared lysate was then applied to a 5 ml HisTrap™ HP column (Cytiva) at a flow rate of 2 ml/min. The column was washed with five column volumes of PBS containing 20 mM imidazole, and the protein was eluted in five column volumes of PBS containing 500 mM imidazole. The N-terminal His_6_-SUMO tag was cleaved by the addition of Ulp1 protease in a 1:50 enzyme to fusion protein ratio (w/w) and 0.14 mM ß-mercaptoethanol, followed by incubation overnight at 4 °C under mild agitation. Subsequently, the reaction mixture was dialyzed against water, lyophilized, and then dissolved in 50 ml of H_2_O Following this, the water-insoluble proteins, contaminants, and aggregates were removed by centrifugation (18,000 rcf for 20 minutes). To cyclize the N-terminal glutamine of CTX to pyroglutamate, the supernatant was acidified by the addition of acetic acid to 5% (v/v) and incubated at 65 °C for 4 h. CTX was then lyophilized again and dissolved once more in a small volume of H_2_O (∼ 20 ml) and subsequently further purified by reversed-phase chromatography on a Zorbax 300SB-C3 column (Agilent, USA) connected to a preparative AZURA-Bio HPLC (KNAUR, Germany) using a linear 5% to 95% acetonitrile gradient in water with 0.1% trifluoroacetic acid. The HPLC fractions were subjected to analysis by MALDI-TOF, and the fractions containing pure CTX with N-terminal pyroglutamate were pooled and lyophilized. Purified CTX was stored at -20 °C for no longer than 6 months.

### Reconstitution of MthK into lipid bilayer nanodiscs

Reconstitution was performed as described previously^8^. Shortly, a 3:1 mixture of POPE:POPG was prepared. The lipids were dissolved in chloroform, mixed, dried under a nitrogen stream, and further dried in a vacuum chamber overnight. The lipid film was dissolved in 20 mM HEPES, 100 mM KCl, pH 7.6, with 2% w/v CHAPS and sonicated until the solution was clear. MthK (monomer) was mixed with MSP1E3D1 and the lipid mixture in molar ratios of 1:2:80. After incubation at room temperature for 1 h, 10 mg Bio-Beads (Bio-Rad™) per 100 µl solution were added. After 3 h, the Bio-Beads were removed, and the same amount of Bio-Beads was added again overnight. The solution was filtered through a 0.22 µm Spin-X centrifugation tube filter (Costar) and loaded on a Superose 6 10/300 column equilibrated with 20 mM HEPES, pH 7.6, 100 mM KCl. The peak containing MthK-loaded nanodiscs was pooled and concentrated up to ∼8 mg/ml using a 100 kDa cut-off Amicon concentrator (Millipore).

### Grid preparation and electron microscopy data collection

MthK-containing nanodiscs were upconcentrated to ∼ 8 mg/ml. Before freezing, the sample was incubated with 200 µM charybdotoxin (CTX) for 30 min at room temperature. Right before freezing, 2 mM fluorinated Fos-choline-8 (Fos8-F, Anatrace) was added, and 3.5 µL sample were applied to a glow-discharged gold grid (UltrAUfoil R1.2/1.3, 300 mesh, Quantifoil). The grid was incubated for 1 s at 22 °C and 100% relative humidity before blotting for 2 s. A Vitrobot Mark IV (FEI, Thermo Fisher Scientific) was used for plunge freezing the grid in liquid ethane. Grids were imaged on a Titan Krios G3i (Thermo Fisher Scientific) operated at 300 kV equipped with a BioQuantum post-column energy filter (slit with 20 eV) and a K3 direct electron detector (Gatan) using EPU software (ThermoFisher Scientific). All micrographs were acquired with aberration-free image shift (AFIS), as dose fractionated movies in super-resolution mode at a nominal magnification of 105kx (resulting in a pixel size of 0.425 Å per pixel on the specimen level) with 64 frames and 0.86 s exposure time. A total dose of 60.76 e^-^/Å^2^ (0.95 e^-^/Å^2^ per frame) and estimated defocus range of -0.6 to -2.6 µm was applied.

### Image processing

Image processing was done in cryoSPARC v4.6.2^33,34^ and Relion-5^53,54^ and is outlined in SI Fig.3. The 20,561 super-resolution movie stacks were motion-corrected and dose-weighted in cryoSPARC Live. The resulting micrographs were analyzed in Relion-5, where CTFFIND4^55^ was used to determine the contrast transfer functions (CTF). Suboptimal micrographs were discarded based on their maximum resolution (8 Å) and relative ice thickness (cut-off value = 8), leaving 15,520 micrographs for further processing. The Laplacian-of-Gaussian picker was used on a subset of 100 micrographs, and the picked particles were extracted four-times binned (pixel size:1.7 Å/pix). After 2D classification, particles showing the complex in high quality were used to train Topaz^56^ on another subset of 100 micrographs. The trained Topaz model was used to pick the whole dataset. Particles were extracted using a box size of 640 pix (pixel size: 0.425 Å/pix), four-times binned (pixel size: 1.7 Å/pix), and submitted to multiple rounds of 2D classification. The resulting 959,199 particles were 3D classified without symmetry into 8 classes using an initial map generated by Relion-5 and low-pass filtered to 60 Å, as a reference. The best class contained 711,385 particles, which were submitted to a 3D classification (with skip-alignment) with a mask around the transmembrane (TM) region and CTX to isolate the bound complex. The resulting 329,943 particles were re-extracted with a pixel size: 0.85 Å/pix. The re-extracted particles were symmetry expanded (C4) and again submitted to a masked 3D classification (with skip-alignment). The particles of the class with the highest resolutions were chosen and duplicates were removed (106,517 particles left). The resulting particles were transferred to cryoSPARC for a local CTF refinement and NU-Refinement (C4), leading to the symmetrical full-length MthK-CTX complex map with a resolution of 3.2 Å. To focus on the non-symmetric CTX, the particles were further analyzed in Relion-5 to yield a C1 map with only one view of the toxin. To enable C1 reconstruction, the gating ring of the 106,517 particles was removed using particle subtraction in Relion-5. The subtracted particles were symmetry-expanded and then used for a guided 3D classification as described in Wu et al. (2024)^31^. A model of the MthK-CTX complex was calculated using AlphaFold3^35^. This model was low-pass filtered to 8 Å resolution and rotated around the symmetry axis in 90° steps. The four resulting models were then used as references for a 3D classification of the subtracted symmetry expanded particles without alignment and a high regularization parameter (T = 60). The class with the best completeness of CTX was chosen for C1 refinement and post-processing, which resulted in a density map with a resolution of 4.1 Å.

### Atomic modeling

Model building for the MthK-CTX complex was started with PDBs of MthK-FL (PDB 5BKI) and CTX (PDB 2CRD). Both structures were docked into the density map using UCSF ChimeraX^57^. The gating ring of MthK-FL was removed to match the cryo-EM density map. ISOLDE^58^ was used to improve the fit to the density map. The resulting complex was refined in real space using PHENIX^59^. Evaluation was performed in MolProbity^60^ and all figures were prepared with ChimeraX^57^.

### Solution NMR-spectroscopy

For solution-state NMR experiments, the lyophilized ^15^N-labeled CTX was dissolved in 500 µl of sample buffer (20 mM Tris, 100 mM KCl, 1.4 mM ß-mercaptoethanol, pH 8.0) The sample was stored for at least 2 h at 4 °C, for the β-mercaptoethanol to slowly evaporate, and 50 µl of D_2_O was added directly before the experiment into the NMR tube. For the spectrum with additional calcium, 10 mM of CaCl_2_ was added directly to the sample. Measurements were performed on a 600 MHz Bruker NMR Spectrometer equipped with a cryo probe. Spectra were measured at 300 K. For the assignment, 3D ^15^N-filtered NOESY and TOCSY experiments were executed. As a fingerprint spectrum, a 2D ^15^N-HSQC spectrum was measured in the beginning and between the 3Ds, verifying that no changes in the sample quality appear over time. Both experiments used a watergate solvent suppression and deuterium for an internal field-frequency lock^61^. The acquisition and processing involved the utilization of Topspin 3.5 (Bruker BioSpin), while data analysis was conducted using CcpNmr 3.2.0^62^. For the chemical shift analysis regarding protein backbone torsion angles, the TALOS+ software^63^ was used.

### Solid-state NMR-spectroscopy

For proton-detected (H)CαNH and (H)CONH 3D experiments, the fully ^13^C,^15^N-labeled, deuterated, and back-exchanged MthK pore domain in proteoliposomes with and without added CTX (unlabeled) was packed into 1.3 mm rotors^52^. Measurements were performed on a 600 MHz Bruker NMR Spectrometer equipped with a 1.3 mm wide bore 3-channel probe (Bruker BioSpin) under 55 kHz magic-angle spinning. Proton-detected 2D cross-polarization (CP) based (H)NH experiments on the ^15^N-labeled, deuterated, and back-exchanged MthK pore domain with and without unlabeled CTX were recorded under 40 kHz magic-angle spinning on an 800 MHz Bruker NMR spectrometer equipped with a 3-channel wide bore probe (Bruker BioSpin) in 1.9 mm rotors. 2D (H)NH spectra based on INEPT transfers between protons and nitrogens for detecting ammonium ions in the ^13^C,^15^N,^2^H-labeled MthK pore domain were conducted on a 900 MHz Bruker spectrometer equipped with a 4-channel standard-bore VTX probe (Bruker BioSpin) in 1.3 mm rotors. Experiments on the ^15^N-labeled CTX bound to the unlabeled MthK pore domain were conducted on the same setup. All experiments took place at a sample temperature of approx. 15 °C, which was determined using the water resonance chemical shift^64^. All spectra were referenced using the lipid peak at 1.21 ppm, because addition of DSS (4,4-dimethyl-4-silapentane-1-sulfonic sodium salt) often used otherwise for referencing could potentially interfere with the ion channel. For the proton detected experiments, MISSISSIPPI water suppression^65^, or a water suppression using a spoil pulse followed by a train of saturation pulses was used^52^. Acquisition and processing, as well as data analysis, were done as described for the solution-state NMR measurements.

### MD Simulations

A simulation system was prepared by CHARMM-GUI^66^ with the CTX and MthK-PD complex structure derived from cryo-EM. All residues were assigned their standard protonation states at pH 7. N- and C-termini were capped with acetyl and methylamide groups, respectively. The complex structure was embedded in a POPE:POPG (3:1) lipid bilayer and solvated in 100 mM KCl. CHARMM36m^67^ or a modified version of CHARMM36m^42^ were used as force field with the TIP3P^68^ water model. The system was energy minimized for 5000 steps, followed by a multistep equilibration in which protein and lipid restraints were gradually reduced over 10 ns. Production simulations used a 2 fs time step, a Parrinello-Rahman barostat^69^ with semi-isotropic pressure control at 1 atm, and a v-rescale thermostat^70^ set to 303.15 K. Nonbonded interactions were cut off at 12 Å with force-switching between 10 and 12 Å, long-range electrostatics were calculated with particle mesh Ewald^71^, and hydrogens were constrained with the LINCS algorithm^72^. For each system, three 500 ns simulations were run using GROMACS 2021^73^. Trajectories were saved every 100 ps and analyzed using MDAnalysis^74^, NumPy^75^, and SciPy^76^ Python libraries, as well as GROMACS analysis tools. To calculate the binding free energy, the gmx_MMPBSA^38^ package was used with the Generalized Born method. For per-residue decomposition analysis, trajectories from three MD runs were concatenated, and residues within 6 Å from both CTX and MthK were selected. Plots were generated using Matplotlib^77^.

