## Supplementary_Information for "Atomic structure and plasticity of the CTX-MthK complex investigated by cryo-EM, NMR, and MD simulations"

[1]Research Unit Molecular Biophysics, Leibniz-Forschungsinstitut für Molekulare Pharmakologie (FMP), Berlin, Germany.
[2]Research Unit Structural Chemistry and Computational Biophysics, Leibniz-Forschungsinstitut für Molekulare Pharmakologie (FMP), Berlin, Germany.
[3]Institute of Chemistry, Technische Universität Berlin, Berlin, Germany.
[4]Core Facility for Cryo Electron Microscopy of the Charité—Universitätsmedizin Berlin at the Max Delbrück Center, Berlin, Germany.
[5]Max Delbrück Center for Molecular Medicine, Technology Platform Cryo-EM, Berlin, Germany.
[6]Core Facility for NMR Spectroscopy, Leibniz-Forschungsinstitut für Molekulare Pharmakologie (FMP), Berlin, Germany.
[7]Leibniz-Forschungsinstitut für Molekulare Pharmakologie (FMP), Berlin, Germany.
[8]Shanghai Institute of Precision Medicine, Ninth People's Hospital, Shanghai Jiao Tong University School of Medicine, Shanghai, China.
[9]Institut für Biologie, Humboldt-Universität zu Berlin, Germany.

#these authors contributed equally.

### Cryo-EM data collection, refinement, and validation statistics

**Table S1.** Cryo-EM data collection, refinement, and validation statistics.

| | MthK-CTX (C4, cryoSPARC) | MthK-CTX (C1) (EMD-56657) (PDB 28NO) |
|---|---|---|
| **Data collection and processing** | | |
| Magnification | 105000 | 105000 |
| Voltage (kV) | 300 | 300 |
| Electron exposure ($e^{-}/Å^2$) | 60.76 | 60.76 |
| Defocus range (µm) | -0.6 to 2.6 | -0.6 to 2.6 |
| Pixel size (Å) | 0.425 (super-resolution) | 0.425 (super-resolution) |
| Symmetry imposed | C4 | C1 |
| Initial particle images (no.) | 3,502,957 | 3,502,957 |
| Final particle images (no.) | 106,517 | 92,302 |
| Map resolution (Å) | 3.2 | 4.1 |
| FSC threshold | 0.143 | 0.143 |
| Map resolution range (Å) | 1.8-9.2 | 3.8-6.5 |
| **Refinement** | | |
| Initial model used (PDB code) | | 5BKI, 2CRD |
| Model resolution (Å) | | 3.7 |
| FSC threshold | | 0.143 |
| Model resolution range (Å) | | |
| Map sharpening *B* factor ($Å^2$) | 109.3 | -159.294 |
| Model composition | | |
| Non-hydrogen atoms | | 3084 |
| Protein residues | | 393 |
| Ligands | | K:1 |
| *B* factors ($Å^2$) | | |
| Protein | | 53.65 |
| Ligand | | 18.46 |

| | |
|---|---|
| R.m.s. deviations | |
| Bond lengths (Å) | 0.003 |
| Bond angles (°) | 0.624 |
| Validation | |
| MolProbity score | 1.03 |
| Clashscore | 0.96 |
| Poor rotamers (%) | 0.3 |
| Ramachandran plot | |
| Favored (%) | 96.34 |
| Allowed (%) | 3.66 |
| Disallowed (%) | 0.0 |

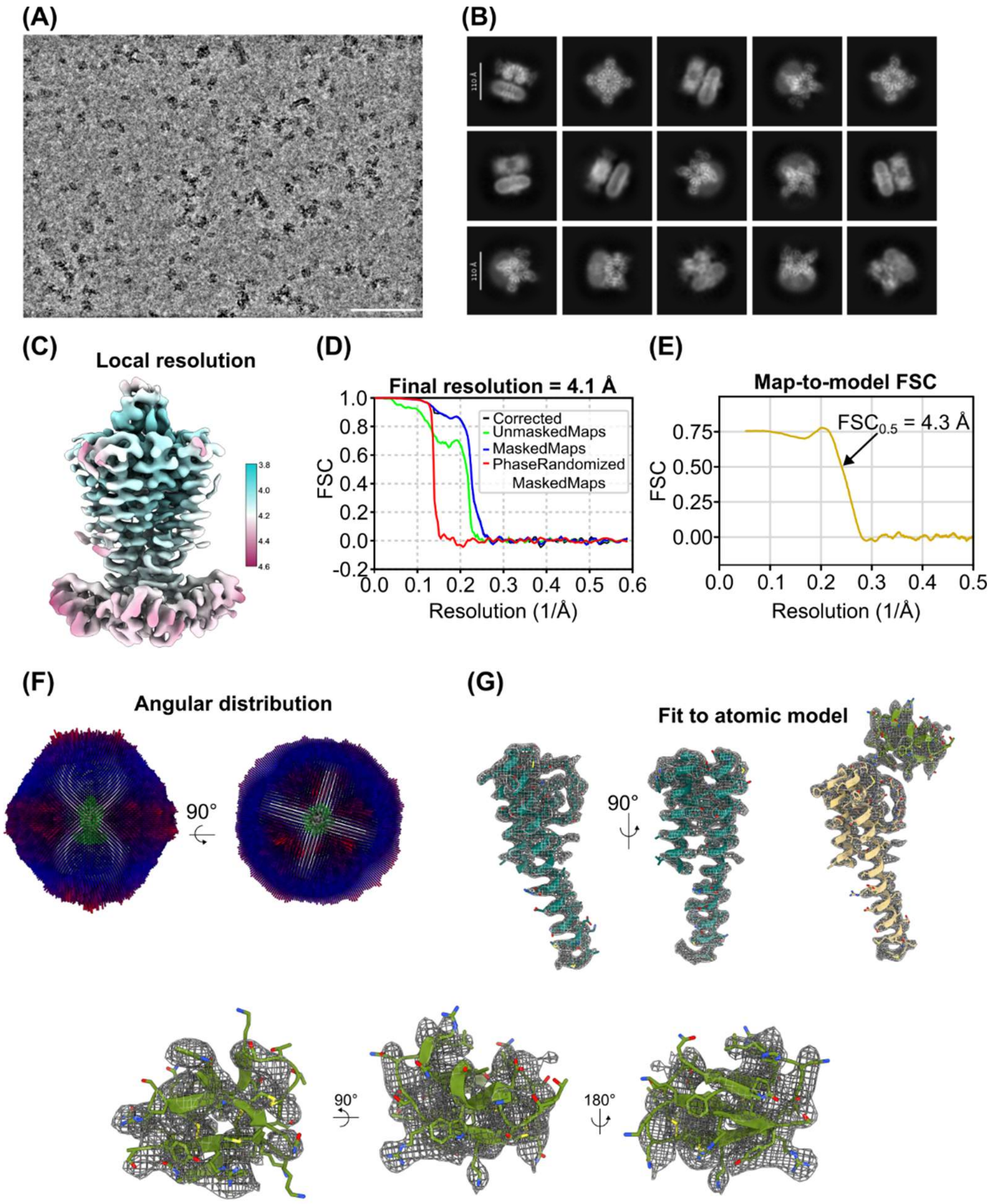

**Figure S1. (A)** Representative cryo-EM micrograph recorded at 300 kV. Scale bar: 100 nm. **(B)** Representative 2D class averages, showing the MthK-CTX complex in lipid nanodiscs from different views. Scale bar: 110 Å. **(C)** Density map colored by local resolution of the final cryo-EM density map of the MthK/CTX complex from 92,302 particles with C1 applied. **(D)** Map-to-model correlation of the MthK/CTX model against the final unsharpened map. **(E)** Fourier shell correlation (FSC) of the final asymmetric MthK-CTX complex. **(F)** Angular distribution of the final particles. **(G)** Fit of the atomic model in selected parts of the density map. Shown is the final post-processed map.

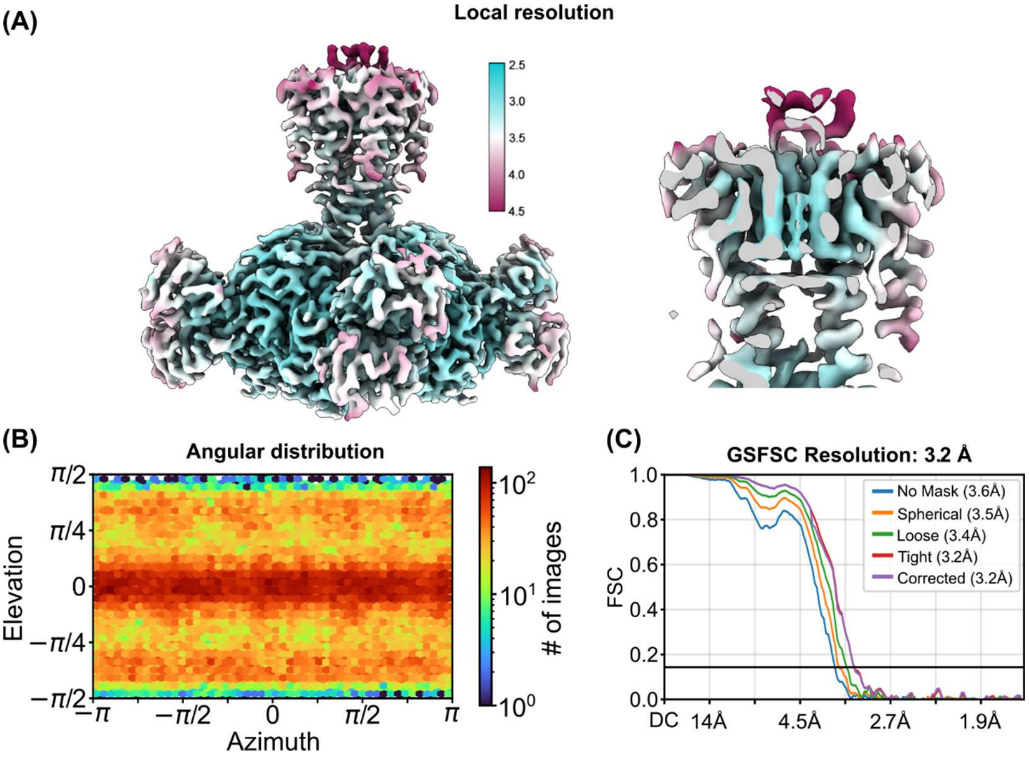

**Figure S2. (A)** Density map colored by local resolution of the full-length MthK-CTX complex from 106,517 particles with C4 applied. **(B)** Angular distribution of the final particles. **(C)** Fourier shell correlation (FSC) of the full-length MthK-CTX complex.

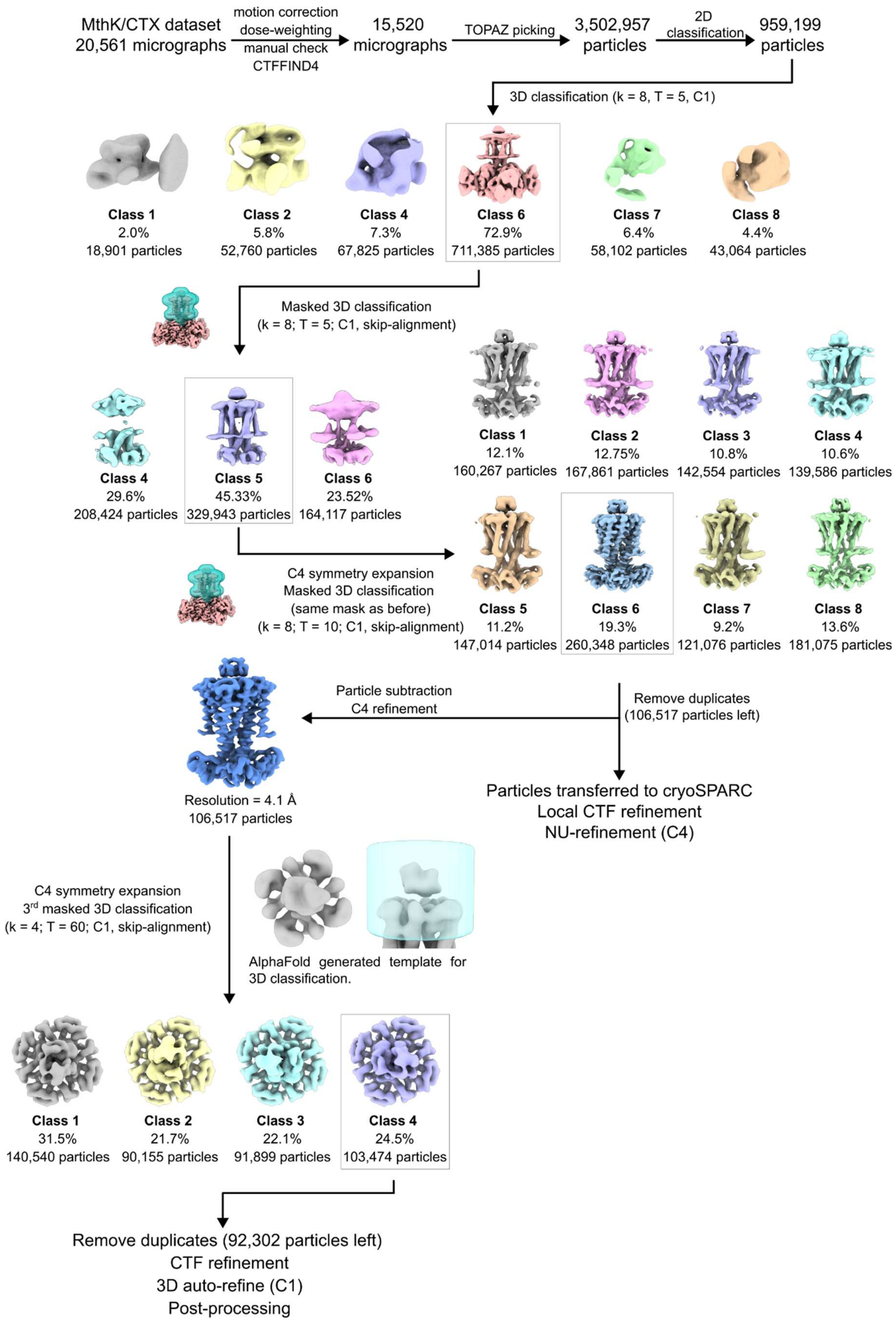
MthK/CTX dataset
20,561 micrographs
motion correction
dose-weighting
manual check
CTFFIND4
15,520
micrographs
TOPAZ picking
3,502,957
particles
2D
classification
959,199
particles
3D classification (k = 8, T = 5, C1)
Class 1
2.0%
18,901 particles
Class 2
5.8%
52,760 particles
Class 4
7.3%
67,825 particles
Class 6
72.9%
711,385 particles
Class 7
6.4%
58,102 particles
Class 8
4.4%
43,064 particles
Masked 3D classification
(k = 8; T = 5; C1, skip-alignment)
Class 4
29.6%
208,424 particles
Class 5
45.33%
329,943 particles
Class 6
23.52%
164,117 particles
Class 1
12.1%
160,267 particles
Class 2
12.75%
167,861 particles
Class 3
10.8%
142,554 particles
Class 4
10.6%
139,586 particles
C4 symmetry expansion
Masked 3D classification
(same mask as before)
(k = 8; T = 10; C1, skip-alignment)
Class 5
11.2%
147,014 particles
Class 6
19.3%
260,348 particles
Class 7
9.2%
121,076 particles
Class 8
13.6%
181,075 particles
Particle subtraction
C4 refinement
Remove duplicates
(106,517 particles left)
Resolution = 4.1 Å
106,517 particles
Particles transferred to cryoSPARC
Local CTF refinement
NU-refinement (C4)
C4 symmetry expansion
3rd masked 3D classification
(k = 4; T = 60; C1, skip-alignment)
AlphaFold generated template for
3D classification.
Class 1
31.5%
140,540 particles
Class 2
21.7%
90,155 particles
Class 3
22.1%
91,899 particles
Class 4
24.5%
103,474 particles
Remove duplicates (92,302 particles left)
CTF refinement
3D auto-refine (C1)
Post-processing

**Figure S3.** SPA data processing scheme applied for the full-length and truncated MthK-CTX complex. Preprocessing (motion correction and dose-weighting) was performed in cryoSPARC. All subsequent steps were carried out in Relion-5, except for the non-uniform refinement of the full-length MthK-CTX complex (C4), which was performed in cryoSPARC.

**Table S2.** Assigned chemical shifts of $^{15}N$-labeled CTX in sample buffer acquired with solution NMR. Equivalent protons are marked with an asterisk (*). The chemical shift assignments were deposited in the BMRB (accession code: 53511).

| | | Chemical shift / ppm | | | | | | |
|---|---|---|---|---|---|---|---|---|
| **Residue Nr.** | **Residue** | **HN** | **N** | **HA** | **HB1** | **HB2** | **HG*** | **additional** |
| 3 | THR | 8.06 | 113.32 | 4.97 | 4.40 | | 1.31 | |
| 4 | ASN | 8.60 | 119.78 | | | | | |
| 5 | VAL | 8.30 | 123.46 | 4.03 | 1.90 | | 1.07 | |
| 6 | SER | 8.69 | 124.08 | 5.15 | 4.01 | | | |
| 7 | CYS | 8.08 | 116.46 | 4.93 | 3.18 | 2.98 | | |
| 8 | THR | 9.67 | 112.47 | 4.84 | 4.53 | | 1.29 | |
| 9 | THR | 8.36 | 114.15 | 4.93 | 4.51 | | 1.29 | |
| 10 | SER | 8.73 | 123.52 | | | | | |
| 11 | LYS | | | | | | | |
| 12 | GLN | 7.37 | 117.40 | 4.14 | | | | |
| 13 | CYS | 7.61 | 113.79 | 4.78 | 3.04 | 2.88 | | |
| 14 | TRP | 7.87 | 125.22 | 4.61 | 3.56 | | | HE 10.02 |
| 15 | SER | | | | | | | |
| 16 | VAL | 7.17 | 123.03 | 3.76 | 2.23 | | 1.26 | 1.03 |
| 17 | CYS | 8.62 | 117.51 | 4.52 | 3.11 | | | |
| 18 | GLU | 8.06 | 124.41 | 3.89 | 2.18 | | 2.40 | HE1 7.36<br>HE2 6.70<br>NE 111.52 |
| 19 | ARG | 7.83 | 118.70 | 4.21 | 2.03 | | 1.77 | HD* 3.32 |
| 20 | LEU | 8.48 | 116.30 | 4.21 | 1.69 | | | HD* 0.92 |
| 21 | HIS | 7.99 | 113.82 | 5.03 | 2.75 | 3.53 | | |
| 22 | ASN | 7.86 | 116.88 | 4.82 | 3.25 | 2.78 | | |

| | | | | | | | | |
|---|---|---|---|---|---|---|---|---|
| 23 | THR | 7.46 | 110.98 | 4.86 | 4.24 | | 1.11 | |
| 24 | SER | 8.29 | 117.85 | 4.81 | 3.88 | | | |
| 25 | ARG | 7.94 | 121.22 | 4.34 | 1.89 | | 1.56 | HD* 3.09 |
| 26 | GLY | 7.62 | 108.69 | 3.86 | 5.23 | | | |
| 27 | LYS | 9.37 | 118.64 | 4.85 | 1.92 | | 1.48 | HD* 2.96 |
| 28 | CYS | 8.85 | 125.07 | 4.85 | 2.76 | | | |
| 29 | MET | 9.00 | 130.49 | 4.89 | 2.20 | 1.92 | 2.55 | |
| 30 | ASN | | | | | | | |
| 31 | LYS | 8.65 | 107.16 | 3.96 | 2.27 | | 1.47 | HD* 1.82 |
| | | | | | | | | HE* 3.11 |
| 32 | LYS | 7.75 | 119.32 | 5.40 | 1.87 | | 1.50 | HE* 3.09 |
| 33 | CYS | 8.60 | 118.19 | 5.25 | 2.68 | 2.99 | | |
| 34 | ARG | 9.68 | 126.94 | 5.00 | 1.76 | | 1.14 | HD* 2.74 |
| 35 | CYS | 8.83 | 123.98 | 5.58 | 2.52 | 3.13 | | |
| 36 | TYR | 8.38 | 122.79 | 4.91 | 2.70 | 3.12 | | |
| 37 | SER | 8.23 | 123.35 | | | | | |

**Table S3.** Assigned chemical shifts of $^{13}C$,$^{15}N$ labeled and deuterated MthK pore domain in $K^+$- and $Ca^{2+}$-containing buffer, obtained from solid-state NMR experiments with and without unlabeled CTX. Assignments obtained from experiments on CTX-washed-in and $NH_4Cl$ buffer samples are marked with * and **, respectively.

| | | | With CTX | Without CTX |
|---|---|---|---|---|
| **Residue Nr.** | **Residue** | **Atom** | **Chemical shift / ppm** | **Chemical shift / ppm** |
| 41 | ILE | C | 176.66 | 176.79 |
| 41 | ILE | CA | | 64.07 |
| 41 | ILE | H | | |
| 41 | ILE | N | | |
| 42 | GLU | C | 177.57 | 177.74 |
| 42 | GLU | CA | 54.94 | 55.04 |
| 42 | GLU | H | 7.20 | 7.21 |
| 42 | GLU | N | 114.51 | 114.30 |
| 43 | GLY | C | 174.81 | 174.78 |
| 43 | GLY | CA | 45.94 | 46.00 |
| 43 | GLY | H | 6.61 | 6.67 |
| 43 | GLY | N | 106.09 | 106.16 |
| 44 | GLU | C | 176.02 | 176.12 |
| 44 | GLU | CA | 53.84 | 53.97 |
| 44 | GLU | H | 5.84 | 5.84 |
| 44 | GLU | N | 116.08 | 116.08 |
| 45 | SER | C | 175.64 | 175.69 |
| 45 | SER | CA | 57.31 | 57.41 |
| 45 | SER | H | 9.19 | 9.34 |
| 45 | SER | N | 117.67 | 117.99 |
| 46 | TRP | C | 177.74 | 177.84 |
| 46 | TRP | CA | 61.41 | 61.42 |

| | | | | |
|---|---|---|---|---|
| 46 | TRP | H | 9.29 | 9.27 |
| 46 | TRP | N | 125.14 | 125.12 |
| 47 | THR | C | 175.85 | 175.78 |
| 47 | THR | CA | 67.09 | 67.33 |
| 47 | THR | H | 8.58 | 8.68 |
| 47 | THR | N | 114.12 | 114.71 |
| 48 | VAL | C | | 177.46 |
| 48 | VAL | CA | 65.93 | 66.03 |
| 48 | VAL | H | 7.59 | 7.60 |
| 48 | VAL | N | 120.92 | 120.70 |
| 59 | THR | C | 172.30** | 171.75 |
| 59 | THR | CA | | |
| 59 | THR | CB | 68.50** | 67.96 |
| 59 | THR | H | - | - |
| 59 | THR | N | - | - |
| 60 | VAL | C | 176.12** | 176.63 |
| 60 | VAL | CA | 65.92** | 66.09 |
| 60 | VAL | H | - | - |
| 60 | VAL | N | - | - |
| 61 | GLY | C | 173.63 | 173.25 |
| 61 | GLY | CA | 47.61** | 46.75 |
| 61 | GLY | H | 7.25* | 6.99 |
| 61 | GLY | N | 101.56* | 99.61 |
| 62 | TYR | C | 177.14 | 178.58 |
| 62 | TYR | CA | 59.01 | 59.19 |
| 62 | TYR | H | 5.73 | 5.69 |

| | | | | |
|---|---|---|---|---|
| 62 | TYR | N | 112.12 | 110.08 |
| 63 | GLY | C | 177.62 | 175.21 |
| 63 | GLY | CA | 45.77 | 44.69 |
| 63 | GLY | H | 9.36 | 9.64 |
| 63 | GLY | N | 99.18 | 100.88 |
| 64 | ASP | C | | 175.54 |
| 64 | ASP | CA | 55.30 | 55.21 |
| 64 | ASP | H | 8.43 | 9.47 |
| 64 | ASP | N | 122.70 | 120.75 |
| 65 | TYR | C | 174.26 | 174.22 |
| 65 | TYR | CA | 56.93 | 57.32 |
| 65 | TYR | H | 7.49 | 7.35 |
| 65 | TYR | N | 115.65 | 115.57 |
| 66 | SER | C | | 169.78 |
| 66 | SER | CA | 56.15 | 56.60 |
| 66 | SER | H | 8.02 | 8.42 |
| 66 | SER | N | 114.69 | 115.40 |
| 67 | PRO | C | 175.82 | 175.86 |
| 67 | PRO | CA | | 62.28 |
| 67 | PRO | H | | |
| 67 | PRO | N | | |
| 68 | SER | C | 174.37 | 174.64 |
| 68 | SER | CA | 58.02 | 58.07 |
| 68 | SER | H | 9.71 | 9.79 |
| 68 | SER | N | 116.19 | 115.85 |
| 69 | THR | C | | 173.20 |

| | | | | |
|---|---|---|---|---|
| 69 | THR | CA | 56.73 | 58.65 |
| 69 | THR | H | 8.39 | 8.57 |
| 69 | THR | N | 115.75 | 115.31 |
| 70 | PRO | C | 178.19 | 178.25 |
| 70 | PRO | CA | | 65.69 |
| 70 | PRO | H | | |
| 70 | PRO | N | | |
| 71 | LEU | C | 178.59 | 178.59 |
| 71 | LEU | CA | 57.98 | 58.08 |
| 71 | LEU | H | 8.52 | 8.62 |
| 71 | LEU | N | 116.20 | 116.42 |
| 72 | GLY | C | 177.86 | 177.90 |
| 72 | GLY | CA | 46.74 | 46.91 |
| 72 | GLY | H | 8.04 | 8.14 |
| 72 | GLY | N | 106.46 | 106.82 |
| 73 | MET | C | 177.50 | 177.77 |
| 73 | MET | CA | 60.46 | 60.66 |
| 73 | MET | H | 8.64 | 8.75 |
| 73 | MET | N | 126.69 | 126.71 |
| 74 | TYR | C | | |
| 74 | TYR | CA | 61.97 | 62.80 |
| 74 | TYR | H | 8.31 | 8.37 |
| 74 | TYR | N | 117.88 | 118.18 |

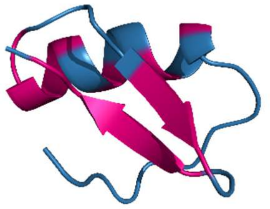

**Figure S4.** TALOS+ prediction mapped on the atomic CTX structure (PDB ID: 2CRD). Pink residues match the secondary structure prediction. Blue residues are predicted to be coil regions or couldn't be unambiguously assigned.

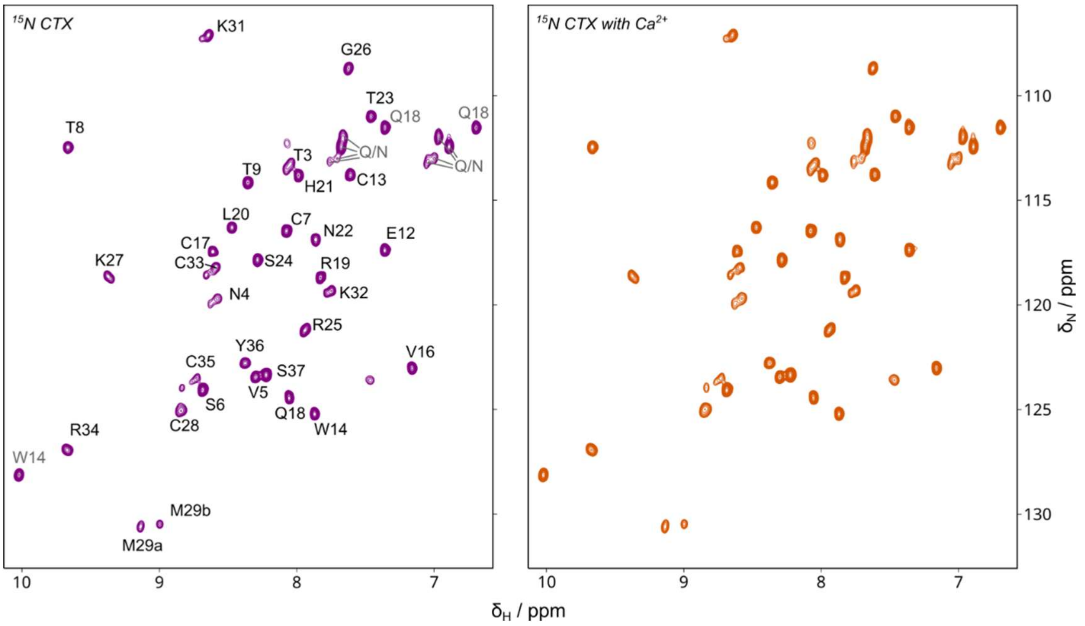

**Figure S5.** $^{15}N$-$^{1}H$ HSQC solution NMR spectra of free CTX in buffer without (left) and with (right) $Ca^{2+}$. Assignments are marked in the left spectrum. The spectra were recorded at 600 MHz at a temperature of 300 K.

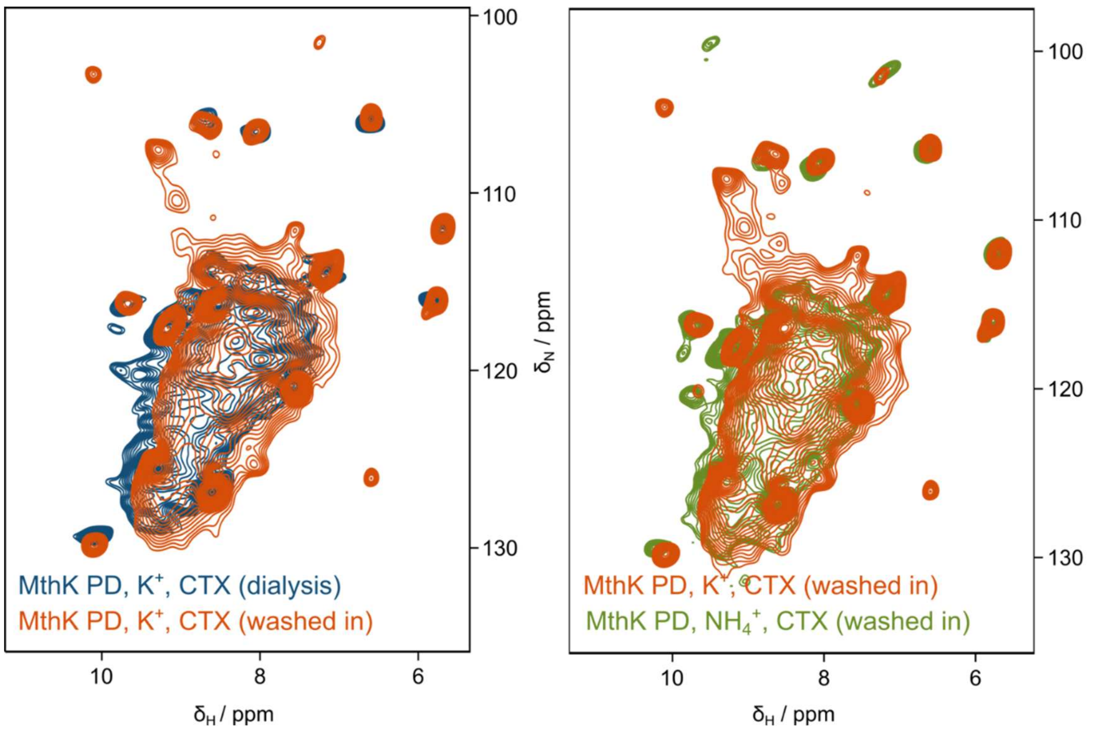

**Figure S6.** (H)NH spectra of MthK-PD. MthK-PD in 100 mM potassium buffer is shown with CTX added during dialysis (blue) and washed in after reconstitution (red). In green, MthK-PD in $^{15}NH_4^+$-buffer and with CTX washed in is shown.

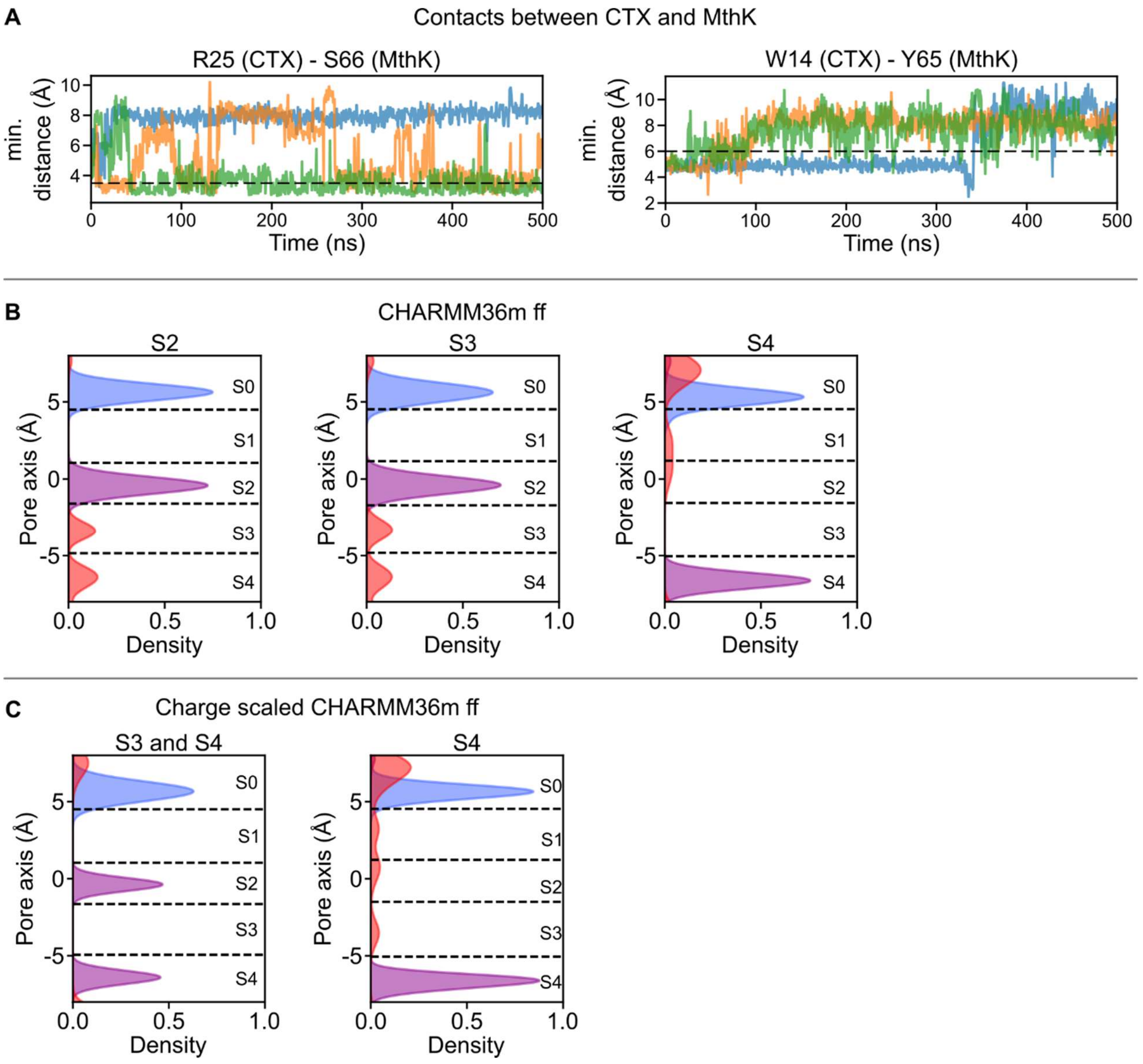

**Figure S7.** Additional data of MD Simulations. (A) The minimum distances between R25/W14 (CTX) and S66/Y65 (MthK) are shown separately, with each line corresponding to one MD run. The dashed black line indicates the distance in the model. (B) The occupancies of K27 (blue) and $K^+$ (purple) in the SF binding sites during MD simulations with the CHARMM36m force field using different initial ion configurations (one ion in either S2, S3, or S4). In all cases, the SF collapsed, with water entering. (C) The occupancies of K27 (blue) and $K^+$ (purple) in the SF binding sites during MD simulations with the charge-scaled CHARMM36m force field using different initial ion configurations (ions in S3 and S4, or only S4). While the latter shows an inactivated SF with water entering, the SF stays stable when two ions are present in S3 and S4 at the beginning. However, the S3 ion moves to S2 as seen in non-charge-scaled MD Simulations.

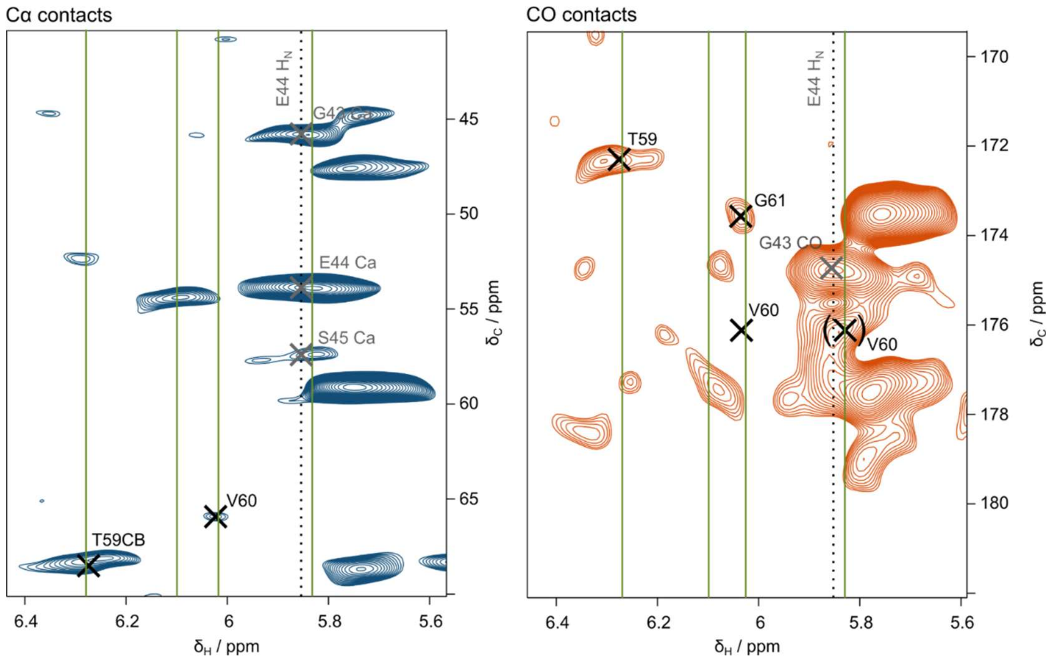

**Figure S8.** (H)CH CP spectra of MthK-PD with washed in CTX, in $^{15}NH_4Cl$-buffer. Experiments were measured with a 7 ms CP mixing time. Contacts between ammonium ions (H-chemical shift marked with green lines) and C-atoms of selectivity-filter residues are marked. Contacts between the backbone HN of E44 and other carbon atoms are marked in grey.
